# Bayesian nonparametric identification of spectrally distinct neural states

**DOI:** 10.64898/2025.12.20.695571

**Authors:** Shoto Yamada, Simon E. Nagel, Xenia Kobeleva, Robert Schmidt

## Abstract

Describing how neural signals change over time is essential for linking fast brain dynamics to underlying cognitive processes. However, this is challenging because neural events can be brief and embedded in dominant broadband aperiodic activity, and may comprise an unknown number of spectrally distinct states. Conventional approaches often apply narrowband band-pass filters to one or a few predefined frequency bands and then use amplitude thresholding to identify oscillatory events, but detection outcomes can be highly sensitive to these choices. Although recent unsupervised alternatives based on hidden Markov models (HMMs) address these limitations, they still require the number of states to be specified in advance and can underfit or overfit when this number is misspecified. We propose a Bayesian nonparametric method that represents temporal dynamics of oscillatory and aperiodic activity as a sequence of discrete states, while inferring an appropriate number of states directly from the data. This method combines time-delay embedding (TDE) with the Dirichlet-process Gaussian mixture model (DP-GMM). TDE augments the signal with time-shifted copies of itself, enabling the DP-GMM to capture local autocovariance structures reflecting underlying spectral features, while the Dirichlet-process prior adapts model complexity by pruning inactive components. We benchmarked the approach against a conventional amplitude thresholding method and the time-delay embedded HMM using single-channel synthetic data designed to mimic neural time series (e.g., EEG, MEG, and local field potentials), with multiple frequency components masked by aperiodic activity. The time-delay embedded DP-GMM not only accurately detects oscillations across multiple frequency bands but also distinguishes between non-oscillatory components with different aperiodic exponents. Applied to a resting-state motor-cortex MEG dataset, the model identified several frequency-selective, short-lived oscillatory states alongside aperiodic states with different spectral components. Overall, this provides an unsupervised framework for discovering multiple spectrally distinct neural states without predefining frequency bands or fixing the number of states.

**Author summary:** Describing complex neural signals can help us link rapid changes in brain activity to underlying brain processes. However, this is challenging because the patterns of neural activity of interest can be brief and difficult to distinguish from noise. Moreover, we typically do not know how many unique neural patterns, or neural states, are present in a recording. To address these challenges, we developed a new unsupervised method that describes a noisy neural signal as a sequence of hidden states and uses the signal itself to estimate how many distinct states are present in the dataset. We tested how accurately our method recovered these hidden neural states in simulated signals and evaluated it using real brain recordings. This approach redefines neural signal analysis by shifting the focus from detecting a specific pattern of neural activity to discovering multiple hidden neural states with distinct patterns of activity. Researchers can then investigate how these states relate to cognitive processes and behavior.

## Introduction

Describing how neural signals change over time is essential for linking fast brain dynamics to underlying cognitive processes. Oscillations in a signal are often of particular interest, as they support frequency-specific coordination and large-scale integration of neural activity [1–4]. Consistent with this view, oscillatory activity has been associated with a broad range of cognitive functions, including, e.g., executive control [5, 6] and memory consolidation [7, 8]. Crucially, many task-relevant oscillations manifest as transient, burst-like events rather than sustained rhythms [9–13], making their reliable identification a nontrivial methodological problem. Such events can be obscured by a dominant broadband aperiodic activity [14], can occur at variable times relative to task events [15], and may appear artificially sustained when brief, temporally jittered episodes are averaged across trials or samples [14, 16].

One conventional approach for identifying oscillatory events is to restrict analysis to one or a few predefined frequency bands via narrowband band-pass filtering, followed by amplitude- or power-thresholding [17–20]. However, the selection of filter and threshold settings can drastically affect detection outcomes [21, 22]. Moreover, if the true oscillation frequency deviates from the filter frequency, the method can falsely detect oscillations in the filtered band, even when no narrowband oscillation is present in the signal [14]. In addition, the threshold directly determines the apparent duration of detected oscillations, and in practice threshold selection often follows arbitrary conventions rather than principled criteria [12]. These issues become even more severe when multiple oscillatory components are present simultaneously. Threshold-based pipelines effectively operate as single-band event detectors and are therefore not well-suited to characterizing multi-frequency structures.

Although a recently developed unsupervised Hidden Markov Model (HMM) approach addresses these issues [23, 24], it still requires pre-specification of the number of states *K*. Typically, researchers fit HMMs with multiple candidate values of *K* and select the preferred model using model-selection criteria such as information criteria or the evidence lower bound (ELBO). This model selection problem remains notoriously challenging [25]. Simulation studies show that commonly used information criteria (e.g., Akaike information criterion; [26], Bayesian Information criterion; [27]) often fail to recover the true number of states, typically favoring overly complex models [25, 28, 29]. For electrophysiological data, the ELBO typically continues to increase with larger *K*, simply favoring models with more states [30, 31]. As a result, practitioners often resort to heuristic choices of *K* guided by the interpretability or reproducibility of their results [23, 24]. Consequently, both formal model-selection criteria and the heuristics leave analyses vulnerable to underfitting (i.e., too few states to capture all the oscillatory states) and overfitting (i.e., fragmenting states into multiple superfluous states).

Motivated by these limitations, we propose a Bayesian nonparametric approach for characterizing temporal dynamics of oscillatory and aperiodic activity as a sequence of discrete states while estimating the number of states. Our method combines time-delay embedding (TDE) [23] with a Dirichlet-process Gaussian mixture model (DP-GMM) [32]. TDE augments the signal with time-shifted versions of itself, enabling the DP-GMM to capture frequency-specific local autocovariance structures. In parallel, the Dirichlet-process prior induces sparsity in the mixture weights, concentrating mass on a small set of data-supported oscillatory states and shrinking redundant components toward near-zero weight.

We evaluated the time-delay embedded DP-GMM on single-channel synthetic time series designed to mimic key properties of neural recordings, including EEG, MEG, and local field potentials, with multiple oscillatory components embedded in broadband aperiodic backgrounds. We compared its performance against two baselines: a conventional amplitude-based thresholding pipeline and the time-delay embedded HMM [23]. We assessed performance across noise levels, oscillatory frequency configurations, and aperiodic spectral exponents, comparing methods with different requirements for prior knowledge of the number of states. We then tested the method on an empirical resting-state MEG dataset, with the number of states selected in a data-driven manner. Across these synthetic and empirical settings, our method identified states characterized by differences in their underlying spectral structures without fixing the number of states a priori, and remained robust to noise and model-order uncertainty. Altogether, the time-delay embedded DP-GMM provides an unsupervised framework for discovering and characterizing multiple spectrally distinct neural states in complex signals without predefining the number of states.

## Materials and methods

### Synthetic data generation

We generated one-dimensional time series that alternate between short-lived oscillatory states and a noise state characterized by a broadband aperiodic activity, following previous studies [23, 33]. A categorical latent state *s*_*n*_ ∈ {1, …, *K*} indexes a noise state (*s*_*n*_ = 1) and *K* − 1 oscillatory states (*s*_*n*_ ≥ 2) for each time point *n*. Signals were constructed by concatenating variable-length segments until a total data point of *N* was reached, truncating the final segment if necessary. We drew an initial state at random from the *K* possible states. If the current state was oscillatory, we switched to the aperiodic state. If it was aperiodic, we drew the next state from the set of oscillatory states.

For each segment of each oscillatory state, we generated a sine wave at a frequency of *f*_sim_. We randomly varied the duration of the oscillatory segment by sampling the number of cycles uniformly from {3, 4, …, 10} . We also scaled the amplitude of each oscillatory state segment by a random factor drawn from | *N* (1, 0.1^2^) | . We finally tapered each segment using a Tukey window with shape parameter 0.25 for a smooth transition between states.

To mimic an aperiodic background activity for a noise state, we first generated a white noise, whose duration was uniformly drawn from 0.2 to 2 seconds. We then filtered the white noise such that its power spectrum decayed as 1*/f*^*β*^, where *β* controls the decay rate. For all simulations we set *β* to 1 (i.e., pink noise) unless stated otherwise.

Unless otherwise noted, we benchmarked the models using 20 simulated time series, each three minutes long and sampled at 250 Hz.

### Signal-to-noise ratio (SNR)

To compute the desired signal-to-noise ratio (SNR), we first high-pass filtered the simulated aperiodic activity at 0.5 Hz to suppress slow drifts, which would otherwise inflate the variance of the signal. Target SNR values were specified in decibels (dB) and converted to a linear power ratio with *γ* = 10^SNR*/*10^. We then scaled the amplitude of each oscillatory state by

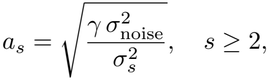

where 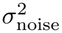 is the power of the noise state (*s* = 1), and 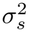 is the power of the oscillatory state (*s* ≥ 2). With this scaling, the power of each state matched the desired SNR relative to the noise power. After scaling all oscillatory states, we added *unfiltered* pink noise to generate the final time series. Thus, the SNR was calibrated using drift-suppressed noise, while the simulated data retained the full 1*/f* structure.

### Performance metric

We quantified the performance of oscillatory state clustering using the adjusted Rand index (ARI) [34]. ARI measures agreement between the estimated and ground-truth state assignments by comparing pairwise state memberships, correcting for chance agreement under a permutation model that preserves the state sizes in each partition [35].

Because ARI is invariant to permutations of state labels, label matching is not required for its calculation. Furthermore, ARI can compare partitions containing different numbers of states, and its chance correction provides a constant expected baseline across different state counts under the permutation model [36]. These properties make ARI suitable for evaluating state recovery even when models overestimate or underestimate the true number of states.

Following the notation of [37], ARI is defined as

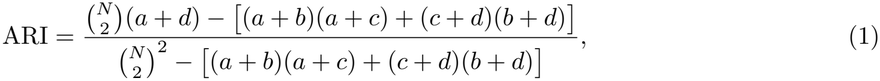

where *N* is the number of time points evaluated. Each unordered pair of time points is classified according to whether its assignments belong to the same state or to different states in each sequence. The four possible outcomes are counted as follows: *a* counts pairs that belong to the same state in both sequences; *b* counts pairs that belong to the same state in the ground truth but to different states in the estimate; *c* counts pairs that belong to different states in the ground truth but to the same state in the estimate; and *d* counts pairs that belong to different states in both sequences. Thus, *a* and *d* represent agreement, whereas *b* and *c* represent disagreement. Together, these counts cover all pairs: 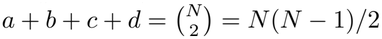. An ARI of 1 indicates perfect agreement, whereas 0 corresponds to the agreement expected by chance under the permutation model. Negative values indicate agreement below this expectation.

For descriptive interpretation, we categorized recovery using previously proposed heuristic thresholds [37]: excellent (ARI ≥ 0.90), good (0.80 ≤ ARI *<* 0.90), moderate (0.65 ≤ ARI *<* 0.80), and poor (ARI *<* 0.65).

### Probabilistic models for clustering spectrally distinct states

#### Feature construction

To detect oscillatory states, we first transformed the raw time series into a feature representation that captures short-range temporal structure by applying TDE (Fig 1). TDE augments the signal with lagged copies of itself, such that the covariance matrix of the embedded vectors reflects the autocovariance of the underlying signal [23].

**Fig 1.**
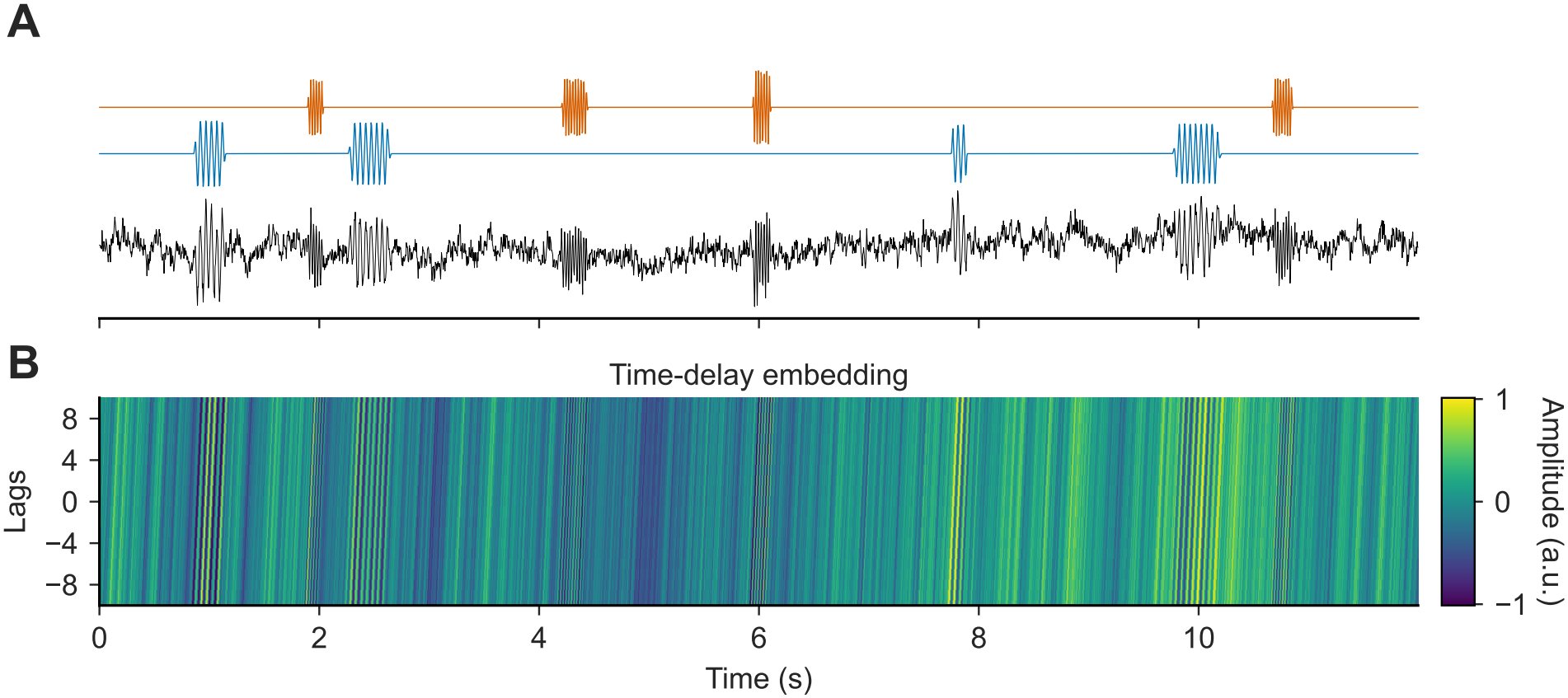
TDE stacks lagged copies of the signal so spectrally distinct states become separable for clustering. (A) Example synthetic signal (black curve) at SNR = 6 dB containing brief 20 Hz (blue) and 40 Hz (orange) oscillatory events. Periods without oscillations consist of pink noise. (B) Corresponding TDE representation with embedding dimension *E* = 21, visualized as lagged copies of the black signal over time. The lag 0 corresponds to the original signal.

The original time series is embedded at a lag of 0. The embedding dimension *E* is a hyperparameter that determines how many lags are included in each embedded vector and thus the effective temporal resolution of the oscillatory structure. *E* was restricted to odd values so that the embedding window was symmetric around the original sample. Consequently, (*E* − 1)*/*2 data points were discarded from each end of the signal.

For both models, clustering performance depended on *E, f*_*s*_, and the target frequency (Fig. S1). In general, clustering performance peaked when the embedding window *w*_*E*_ ≈ *E/f*_*s*_ spanned approximately 2.5 cycles at the target frequency. We therefore set the embedding dimension using the cycle-based heuristic

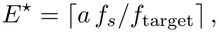

with *a* = 2.5 for both models, where ⌈°⌉ denotes the ceiling operator and *f*_target_ is the target frequency. For example, for *f*_target_ = 30 Hz and *f*_*s*_ = 250 Hz, this gives *E*^*⋆*^ = ⌈2.5 × 250*/*30⌉ = 21. We used this pragmatic, empirically guided heuristic to set the embedding dimension in subsequent analyses.

In addition, we standardized the data and then applied TDE to obtain embedded observations, which served as input to both DP-GMM and the HMM.

#### DP-GMM

The DP-GMM is a Bayesian nonparametric extension of the Gaussian mixture model that allows the data to determine how many mixture components (i.e., states) are effectively used [38]. The DP prior allows for an unbounded number of potential components, and it assigns non-zero weight to only a limited number of them [32]. For computation, we used a truncated stick-breaking approximation with an upper bound *K*_upper_ on the number of states. This provides a finite representation while retaining the DP prior behavior that unused components receive near-zero posterior weights [39].

Stick-breaking generates mixture weights recursively. For each *k* = 1, …, *K*_upper_, we first sample *v*_*k*_ ∼ Beta(1, *α*). Starting from a stick of length 1, at step *k* we break off a fraction *v*_*k*_ of the remaining stick and assign that piece as the *k*th weight *w*_*k*_ (with the remainder passed to the next step). That is, the resulting weights 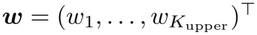 are a deterministic function of 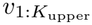 . We denote this construction compactly as

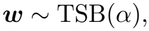

where TSB(*α*) is the truncated stick-breaking prior at *K*_upper_ components. Smaller values of *α* imply faster decay of the expected weights and hence sparser mixtures. We placed a Half-Cauchy(*β*_*α*_ = 1) hyper prior on the concentration parameter *α*, allowing it to be learned from the data.

We modeled observations with a zero-mean multivariate normal distribution,

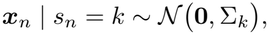

where *s*_*n*_ ∈ {1, …, *K*_upper_} is the latent state at a time point *n* and Σ_*k*_ is the state-specific covariance matrix (treated as a random variable). For computation, we parameterized each covariance matrix via a scale-correlation decomposition,

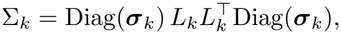

where ***σ***_*k*_ ∈ ℝ^*d*^ is the vector of marginal standard deviations, *L*_*k*_ is the Cholesky factor of a correlation matrix, and *d* is the data dimension (i.e., embedding dimension). This decomposition is a flexible alternative to inverse-Wishart priors for covariance matrices, particularly effective in moderate to high dimensions [40]. We placed independent Half-Cauchy(*β*_*σ*_ = 1) priors on the scale parameters *σ*_*k,j*_ and an LKJCholesky(*η* = 1) prior on *L*_*k*_; for *η* = 1, the LKJ prior is uniform over the space of valid correlation matrices [41]. Together, these priors induce *p*(Σ_*k*_).

Collecting all unobserved random variables into

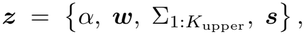

the joint distribution factorizes as

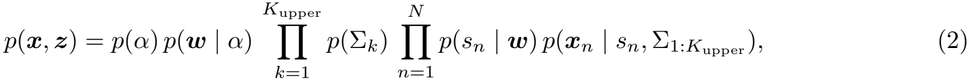

where *N* denotes the total number of data points, *p*(*s*_*n*_ | ***w***) is a categorical distribution with probabilities ***w***, and 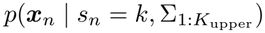 is the Gaussian observation model defined above. Fig 2A provides the graphical representation of the DP-GMM.

**Fig 2.**
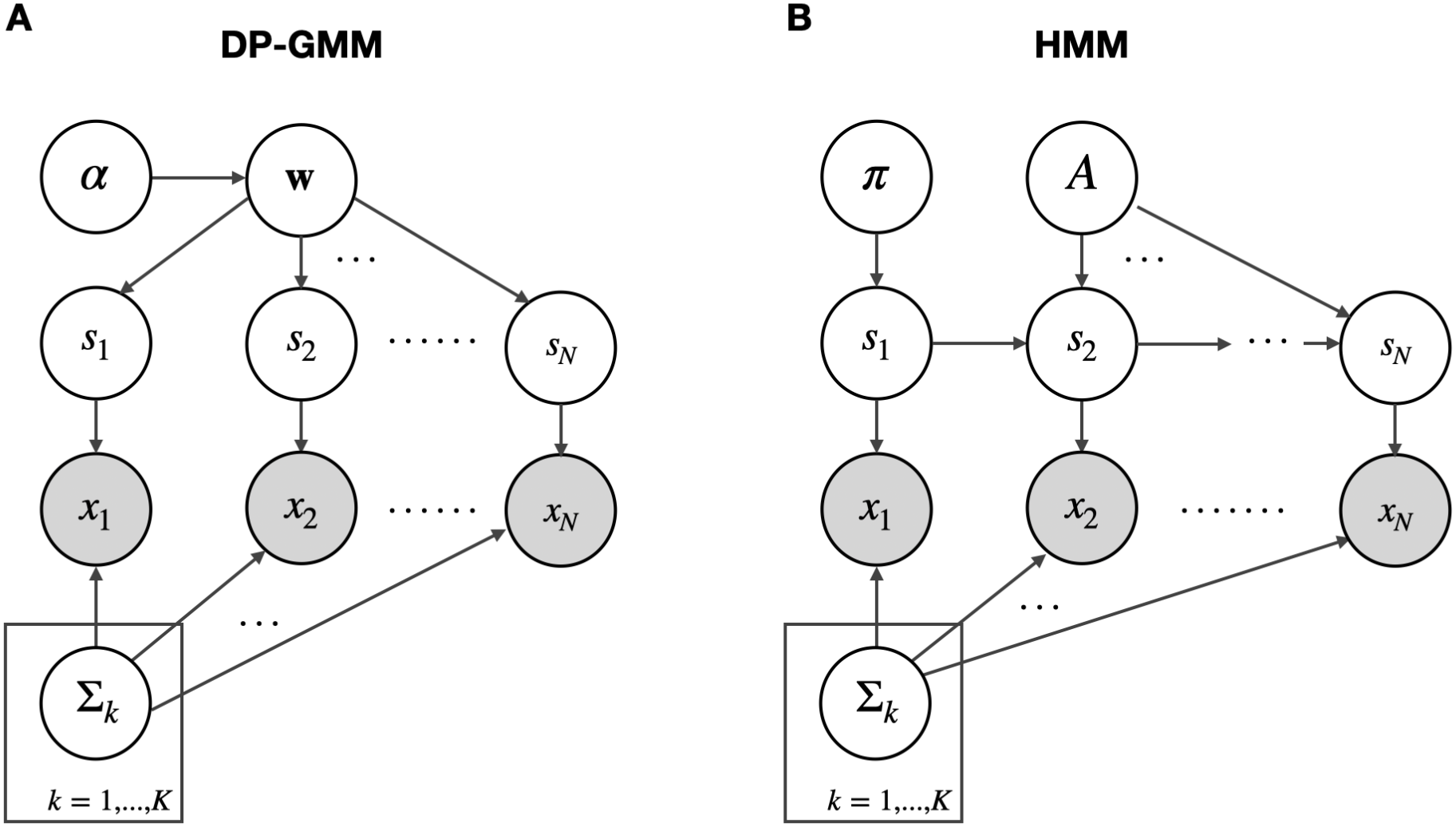
Model overview: DP-GMM and HMM share the same Gaussian emission model but differ in latent-state dynamics. (A) DP-GMM: a concentration parameter *α* induces mixture weights ***w*** (truncated stick-breaking); latent states *s*_1:*N*_ are i.i.d. with class probabilities ***w***. Embedded observations follow ***x***_*n*_ | *s*_*n*_ = *k* ∼ *N* (**0**, Σ_*k*_), where Σ_*k*_ is the state-specific covariance of the time-delay embedded vectors.(B) HMM: an initial distribution ***π*** and transition matrix *A* generate a first-order Markov state sequence *s*_1:*N*_ ; observations follow the same Gaussian emission model. Shaded nodes denote observed variables and plates indicate replication over states *k* = 1, …, *K*.

Unless otherwise stated, we set the upper bound on the number of states to *K*_upper_ = ⌈ln *N*⌉, motivated by the fact that the expected number of occupied clusters under a DP prior grows on the order of ln *N* [42]. Since *K*_upper_ is only an upper bound, the DP prior typically assigns negligible posterior weight to many components. We first retained the smallest set of components with the largest posterior mixture weights that collectively accounted for at least 99% of the total mixture weight across the *K*_upper_ components, treating the remaining components as inactive.

Among the retained components, we further excluded states whose covariance matrices exhibited nonuniform variances across embedding dimensions (i.e., unequal diagonal elements). Typical examples were transitional states at the boundaries of dominant state segments, which we refer to as edge states (Fig. S2). We observed these edge states almost exclusively at high SNRs. These states were very short and had little effect on clustering accuracy, but removing them made the inferred states easier to interpret. We quantified the non-uniformity of variances using the coefficient of variation (CV) of the diagonal elements of each covariance matrix and excluded states with CV values greater than 0.1. We defined the number of active states as the number of remaining components and normalized their weights to sum to one. Using the fitted DP-GMM parameters for these active states, we estimated the state sequence by assigning each time point to the most likely state via maximum a posteriori assignment.

Sensitivity analyses showed that the model’s performance varied little across the tested values of the prior hyperparameters *β*_*α*_, *β*_*σ*_, and *η* (Fig. S3). Clustering accuracy also varied little across much of the tested range of CV thresholds (Fig. S4). Thresholds below approximately 0.05 tended to exclude components corresponding to true states and therefore reduce accuracy, whereas higher thresholds retained additional states at high SNRs. These results supported the threshold of 0.1 as a practical choice for reducing excess states while maintaining clustering accuracy.

#### HMM

The HMM represents a time series as a sequence of *K* discrete latent states with Markovian temporal dynamics [43, 44]. Let *s*_*n*_ ∈ {1, …, *K*} denote the latent state at time point *n*, and ***x***_*n*_ ∈ ℝ^*d*^ the corresponding observation. Conditioned on the current state, observations are generated independently from a state-specific emission model. We used the same zero-mean multivariate normal emission model, and the same covariance prior *p*(Σ_*k*_) as in the DP-GMM (see <u>DP-GMM</u>for details).

The latent state sequence follows a first-order Markov chain governed by an initial state distribution ***π*** and a transition matrix *A*, where *A*_*ij*_ = *p*(*s*_*n*_ = *j* | *s*_*n−*1_ = *i*). Specifically,

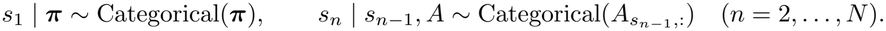

We placed symmetric Dirichlet priors on ***π*** and on each row of *A*,

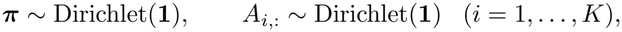

where **1** denotes the all-ones vector of length *K*.

Collecting all latent variables and parameters into

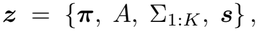

the joint distribution factorizes as

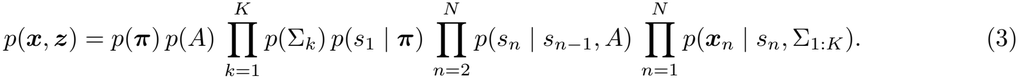

Fig 2B provides the graphical representation of the HMM.

After fitting the model, we decoded a latent state sequence using the Viterbi algorithm [45, 46], which returns the maximum a posteriori state path under the fitted HMM parameters,

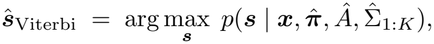

where 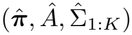 denotes the inferred parameter values.

#### Stochastic variational inference

Given a probabilistic model specified by a joint distribution *p*(*x, z*), Bayesian inference aims to compute the posterior distribution of all unobserved random variables given the observations,

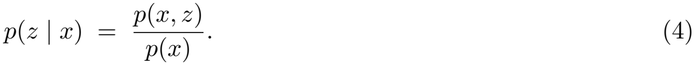

Here, *z* denotes all latent variables in the model, including continuous parameters and discrete latent states. For the models considered here, exact posterior inference is analytically intractable due to integrals that cannot be evaluated in closed form. Moreover, empirical neural time series are typically large, requiring inference methods that scale to big datasets. We therefore used stochastic variational inference (SVI) to obtain a fast and scalable approximation of the posterior via stochastic gradient optimization [47, 48].

SVI approximates *p*(*z* | *x*) with a tractable variational distribution *q*(*z*; *ϕ*), parameterized by *ϕ*, by maximizing the evidence lower bound (ELBO) [49, 50]]:

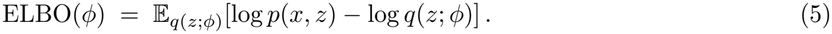

For optimization, we minimized the negative ELBO (NELBO), defined as NELBO(*ϕ*) = − ELBO(*ϕ*); minimizing NELBO(*ϕ*) is equivalent to maximizing ELBO(*ϕ*). Because this expectation is generally intractable, we optimized a Monte Carlo estimate of the NELBO. Concretely, at each optimization step, we drew *L* samples (particles) *z*^(1)^, …, *z*^(*L*)^ ∼ *q*(*z*; *ϕ*) and approximated the NELBO and its gradients by averaging over these samples; increasing *L* reduces estimator variance at a higher computational cost.

We implemented our models and inference in Python using the probabilistic programming libraries NumPyro [51] (for the DP-GMM) and Pyro [52] (for the HMM). Preliminary benchmarks of our implementation showed that NumPyro yielded shorter runtime for the DP-GMM; therefore, we used NumPyro for this model. For the continuous latent variables, we used an automatically constructed mean-field Normal variational family (i.e., a diagonal-covariance Gaussian) in both frameworks. Because both models include discrete latent state variables, we marginalized these variables using parallel enumeration, enabling automatic differentiation for the remaining continuous variables and improving optimization stability [53, 54].

For the DP-GMM, we ran SVI with *L* = 10 Monte Carlo particles, minibatches of size 2^10^ observations sampled uniformly from individual time points, a learning rate of 5 × 10^*−*2^, and 3000 training epochs. For the HMM, we optimized randomly sampled subsequences of length 1000 using minibatches of 2^4^ subsequences, one Monte Carlo particle (*L* = 1), a learning rate of 5 × 10^*−*2^, and 3000 training epochs. These model-specific settings were chosen to obtain stable convergence within feasible runtime and memory constraints, while accounting for differences in minibatching and latent-variable structure between the DP-GMM and HMM. On a MacBook Pro with an Apple M4 Pro chip (14-core CPU, 24 GB of unified memory), wall-clock fitting times were 43.6 ± 0.7 s for the DP-GMM and 226 ± 1.6 s for the HMM (mean ± SD across seven runs per model). Both models were fitted to a three-minute simulated signal sampled at 250 Hz, with the DP-GMM upper bound states and the number of HMM states both set to 11.

To reduce sensitivity to initialization, we fit both models with 10 random initializations. For each fitted run, we estimated the final NELBO using 100 Monte Carlo samples from *q*(*z*; *ϕ*) and retained the run with the lowest final NELBO for subsequent analyses. We summarized each continuous latent variable by the median of its approximate posterior distribution under *q*(*z*; *ϕ*).

### Thresholding method

Among many variants of thresholding methods [33], we used the one with two amplitude thresholds: a main detection threshold and a lower secondary threshold for defining event boundaries [55, 56]. Given a band-pass filtered signal centered at the target frequency, the detection algorithm proceeds in three steps. First, it computes the analytic signal via the Hilbert transform and takes its magnitude as the amplitude envelope. Second, candidate oscillatory events are identified as periods during which the amplitude envelope exceeds the main detection threshold. Third, for each candidate event, it defines its event onset and offset as the time before and after the amplitude envelope falls below the secondary threshold. These steps yield a binary time series that indicates the presence or absence of oscillatory events at each time point.

In the simulation experiments, the band-pass filter was centered at the known ground-truth frequency *f*_*sim*_. We used a Gaussian filter with a full-width at half maximum of 5 Hz around *f*_*sim*_. Threshold parameters were selected in a data-driven manner. We obtained a binary burst-indicator time series from the detection algorithm and selected the threshold pair by maximizing the Pearson correlation between this indicator and the amplitude envelope of the band-pass filtered signal, a pragmatic criterion used in prior work for tuning burst-detection parameters [17, 18].

To extend the thresholding method to scenarios with multiple oscillatory states, each target frequency band was processed individually, with thresholds optimized separately for each band, yielding a binary time series per band. These per-band detections were then combined into a single categorical label per time point. Samples with no detections were labeled as a noise state, whereas samples with one or more detections were assigned to an oscillatory state based on the highest amplitude envelope of the corresponding band.

### Post-hoc *k*-nearest-neighbor imputation

Both the DP-GMM and the HMM occasionally produced brief, fragmented state activations (Fig. S5). This observation motivated a simple post-hoc *k*-nearest-neighbor (*k*-NN) imputation procedure. Contiguous state segments containing fewer than *k* samples were flagged as spurious, and their labels were marked as missing. Missing labels were then processed in temporal order, with each label replaced by the most frequent available state label within a centered window with an initial radius of *r* = *k/*2 samples. At the beginning and end of the recording, the search window included only the available samples.

To select the imputation window duration for each model, we performed a sensitivity analysis for the 20 Hz and 40 Hz simulation condition at a sampling frequency of 250 Hz, varying both SNR and imputation window duration (Fig. S6). Imputation was particularly beneficial at relatively low SNRs (e.g., − 2 to 2 dB). Although brief state fragments typically occurred at similar onset times under both models, they tended to have shorter durations under the DP-GMM than under the HMM (Fig. S5). Consistent with this difference, the DP-GMM benefited from shorter imputation windows, and we selected 30 ms as a conservative default. This window duration was insufficient to improve clustering accuracy with the HMM, so we selected a longer default window of 70 ms. Although longer windows could further improve clustering accuracy, we chose 70 ms to keep the imputation threshold just below the minimum simulated burst duration of 75 ms, corresponding to three cycles at 40 Hz.

Although imputation had little effect on the median ARI for thresholding (Fig. S7), we used the same 30-ms imputation window for thresholding and the DP-GMM for consistency.

### CTF rest MEG dataset

We applied the time-delay embedded DP-GMM to an empirical, publicly available dataset to evaluate whether it could discover neural states that differ in spectral content. We analyzed data from 65 healthy participants recorded on a 275-channel CTF system (5 min per subject) at Nottingham University, UK, as part of the MEGUK partnership [57]. We used the preprocessed parcel-wise time series released by Gohil et al [24]. In brief, 0.5-125 Hz band-pass and 50/100 Hz notch filters were applied; the data were downsampled to 250 Hz; ocular and cardiac artifacts were removed via automated ICA using EOG/ECG correlations; and the data were finally downsampled to 100 Hz. We selected a single parcel in the left motor cortex and temporally concatenated the corresponding time series from all 65 participants to form a single continuous signal, following prior work using the same strategy to enable group-level state inference from resting-state MEG [24].

## Results

### The DP-GMM maintains high clustering accuracy even when the true number of states is unknown

Reliable and practically useful clustering of oscillatory states must be robust to noise and applicable to signals containing multiple oscillatory states. In practice, the true number of states is unknown. We therefore evaluated DP-GMM clustering performance across a series of benchmark scenarios, treating the number of states as unknown. We specified only an upper bound of *K*_upper_ = 11, allowing the model to prune redundant states and select the final number of states automatically.

Unless stated otherwise, performance is reported as the median ARI across 20 simulated samples per condition. Single ARI values refer to the representative example shown in the corresponding figure panel. Random seeds differed across 20 independent simulations but were held fixed across conditions within each sample.

### The DP-GMM reliably recovers oscillatory states

We benchmarked the DP-GMM against the HMM and the thresholding method across a range of SNRs and frequency conditions. Here, we generated 20 independent three-minute simulations with 3 states (two oscillatory states and one noise state) at *f*_*s*_ = 250 Hz with SNR ∈ {− 10, − 8, …, 10} dB and six frequency conditions (10,40), (15,40), (20,40), (25,40), (30,40), (35,40) Hz (Fig. S8). We fitted both models using 10 initializations each to compute a single ARI.

For the HMM, we fixed the number of states at the ground-truth value (*K* = 3), assuming that the true number of states was known. The thresholding method assumed knowledge of both the true number of oscillatory states and the true target frequencies. For each target frequency, we applied a separate Gaussian band-pass filter centered on its ground-truth value (see Thresholding method for algorithm details). These assumptions placed both the HMM and the thresholding method in a best-case configuration.

The embedding dimension was determined using the empirically guided, cycle-based heuristic (see Methods). For each pair of oscillatory states, we defined *f*_target_ as the mean of the two state frequencies and selected the embedding dimension so that the embedding window spanned approximately 2.5 cycles for both the DP-GMM and the HMM. This yielded 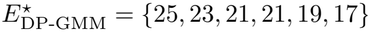 for the 10 and 40, 15 and 40, 20 and 40, 25 and 40, 30 and 40, and 35 and 40 Hz frequency pairs, respectively.

In the representative 20 and 40 Hz simulation at SNR = 2 dB, the DP-GMM (ARI = 0.88) outperformed both the HMM (ARI = 0.75) and the thresholding method (ARI = 0.85), without requiring the true number of states to be specified (Fig. 3A). In this frequency condition, the DP-GMM consistently achieved a higher median ARI than the other methods at SNR ≥ 2 dB (Fig. 3B). Using the descriptive ARI categories defined in the Methods, recovery was excellent at SNR ≥ 4 dB (median ARI ≥ 0.92) and good at SNRs of 0 and 2 dB (median ARIs of 0.82 and 0.88, respectively). At SNR = − 2 dB, recovery remained moderate (median ARI = 0.72). Although omitting post-hoc imputation reduced ARI by approximately 0.10, the DP-GMM still achieved a higher ARI than the HMM overall and than the thresholding method at SNR ≥ 4 dB. This observed advantage of the DP-GMM was generalizable to the most of the other frequency conditions (Fig. 3C). Even in the 35 and 40 Hz condition, where the two frequencies differed by only 5 Hz, the DP-GMM achieved a median ARI ≥ 0.85 at SNR ≥ 0 dB, indicating that it can distinguish states with closely spaced oscillatory frequencies.For the most widely separated frequency pair (10/40 Hz), however, the DP-GMM showed poorer recovery at low to moderate SNRs than for the other frequency pairs. At 0 dB, its median ARI after imputation was 0.57 for the 10/40-Hz pair (poor recovery), compared with 0.82 for the 20/40-Hz pair (good recovery).

**Fig 3.**
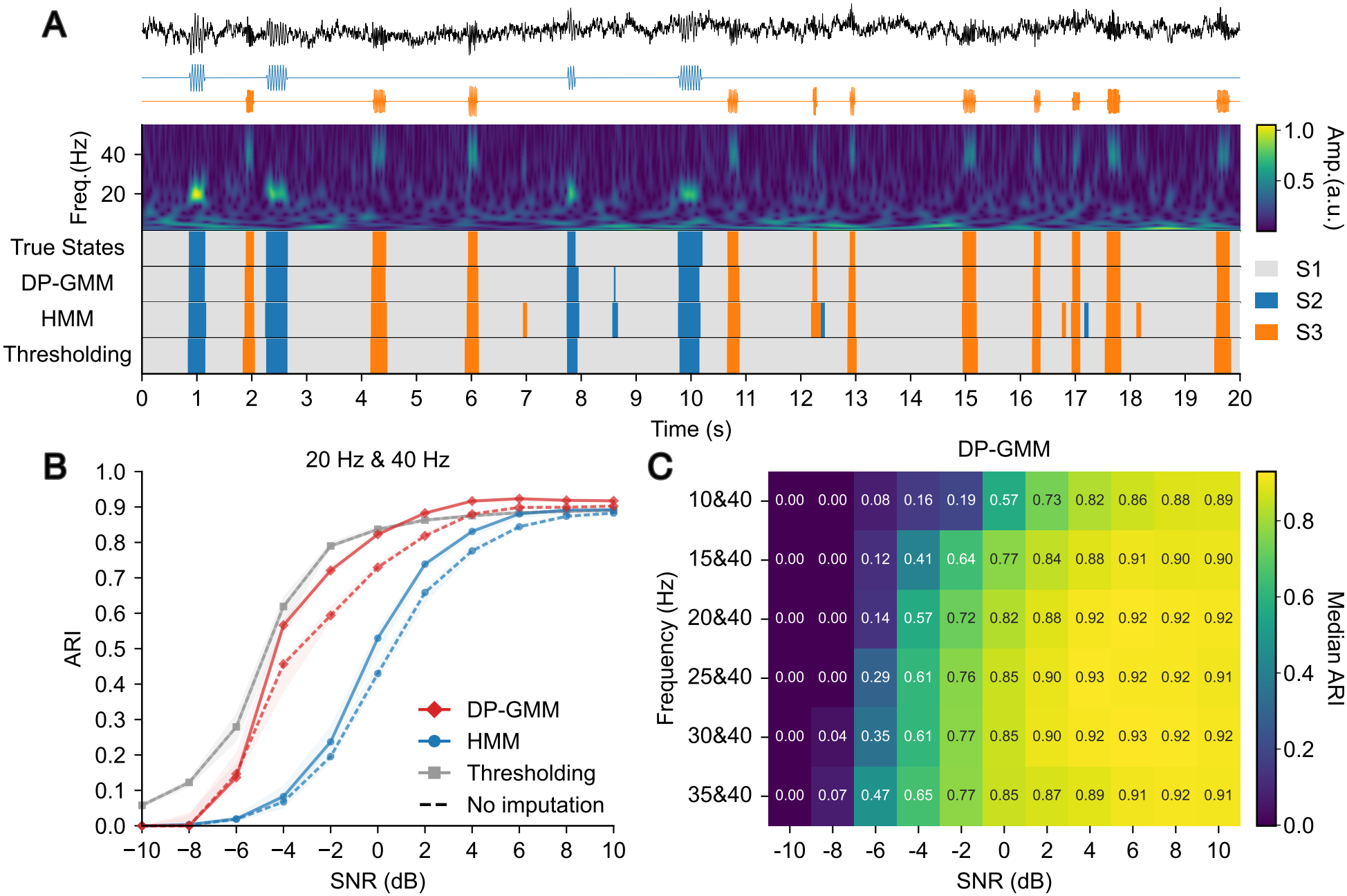
The DP-GMM accurately identifies oscillatory states in noisy signals. (A) Example raw trace from a three-state synthetic signal containing 20 Hz (blue) and 40 Hz (orange) oscillations at SNR = 2 dB, shown with wavelet amplitude (a.u.), the ground-truth state sequence, and the estimated oscillatory states after *k*-NN imputation. (B) Median ARI as a function of SNR for the 20 and 40 Hz condition (solid: with imputation; dashed: without imputation). The shaded area represents the interquartile range. (C) DP-GMM median ARI after imputation across SNRs and frequency-pair conditions.

The HMM was sensitive to noise. With imputation, it first achieved good recovery (median ARI ≥ 0.80) at 4 dB for several frequency pairs (Fig. S7). Without imputation, most pairs required 6 dB. At low to moderate SNRs, the HMM struggled with both the most widely separated pair (10/40 Hz) and the two closest pairs (30/40 and 35/40 Hz). For the 35/40-Hz pair in particular, recovery remained poor even at 10 dB, with median ARIs of 0.28 before imputation and 0.27 after imputation.

Thresholding performance varied less across frequency pairs at a given SNR than did the performance of the DP-GMM and the HMM (Fig. S7). Thresholding outperformed both models for the 10/40-, 15/40-, and 20/40-Hz pairs at all tested SNRs up to and including 0 dB, although all three methods showed poor recovery for these pairs at SNRs of − 4 dB or lower (median ARI *<* 0.65).

In terms of state-count estimation, the DP-GMM correctly estimated the true number of states (i.e., three states) across a wide range of SNRs and frequency conditions (Fig. S9). Although the number of states estimated before CV filtering increased with SNR at SNR ≥ 4 dB because of edge states, the CV-based criterion (CV *<* 0.1) suppressed these states and recovered the correct number at SNR ≥ 2 dB. At very low SNRs (e.g., SNR ≤ − 6 dB), the DP-GMM underestimated the number of states. At SNR ≤ − 8 dB, it returned a single noise state, with no separate oscillatory states identified (Fig. S9).

Overall, the DP-GMM accurately distinguished multiple oscillatory states in noisy signals, including states with closely spaced frequencies, and often recovered the true number of states. It outperformed the HMM across most conditions, both with and without imputation. After imputation, it also generally outperformed the thresholding method at moderate noise levels. These results show that the DP-GMM can achieve high clustering accuracy without requiring the true number of states to be specified in advance.

### The DP-GMM reduces the risks associated with misspecifying the number of states

In this section, we demonstrate the advantage of the DP-GMM over the HMM from another perspective, showing its robustness to misspecification of the number of states. For this purpose, we set the upper bound for the DP-GMM to 11 states and fixed the number of HMM states at 11, deliberately overparameterizing the HMM. For both models, we used the same embedding dimension (*E* = 21), chosen based on two cycles at the mean target frequency of 30 Hz.

The DP-GMM accurately clustered the oscillatory states (ARI = 0.83, Fig. 4A) and correctly retained three active states (Fig. 4B). In contrast, the overparameterized HMM tended to use all 11 states, splitting the noise state into many extremely brief fragments (ARI = 0.07, Fig. 4A). Its stationary distribution was also nearly uniform across all 11 states, indicating similar expected long-run occupancy probabilities for each state (Fig. 4C). Because these fragments dominated local neighborhoods in the HMM state sequence, *k*-NN imputation would not be expected to recover the true number of states, whereas DP-GMM clustering performance could be further improved by imputing brief and rare fragments.

**Fig 4.**
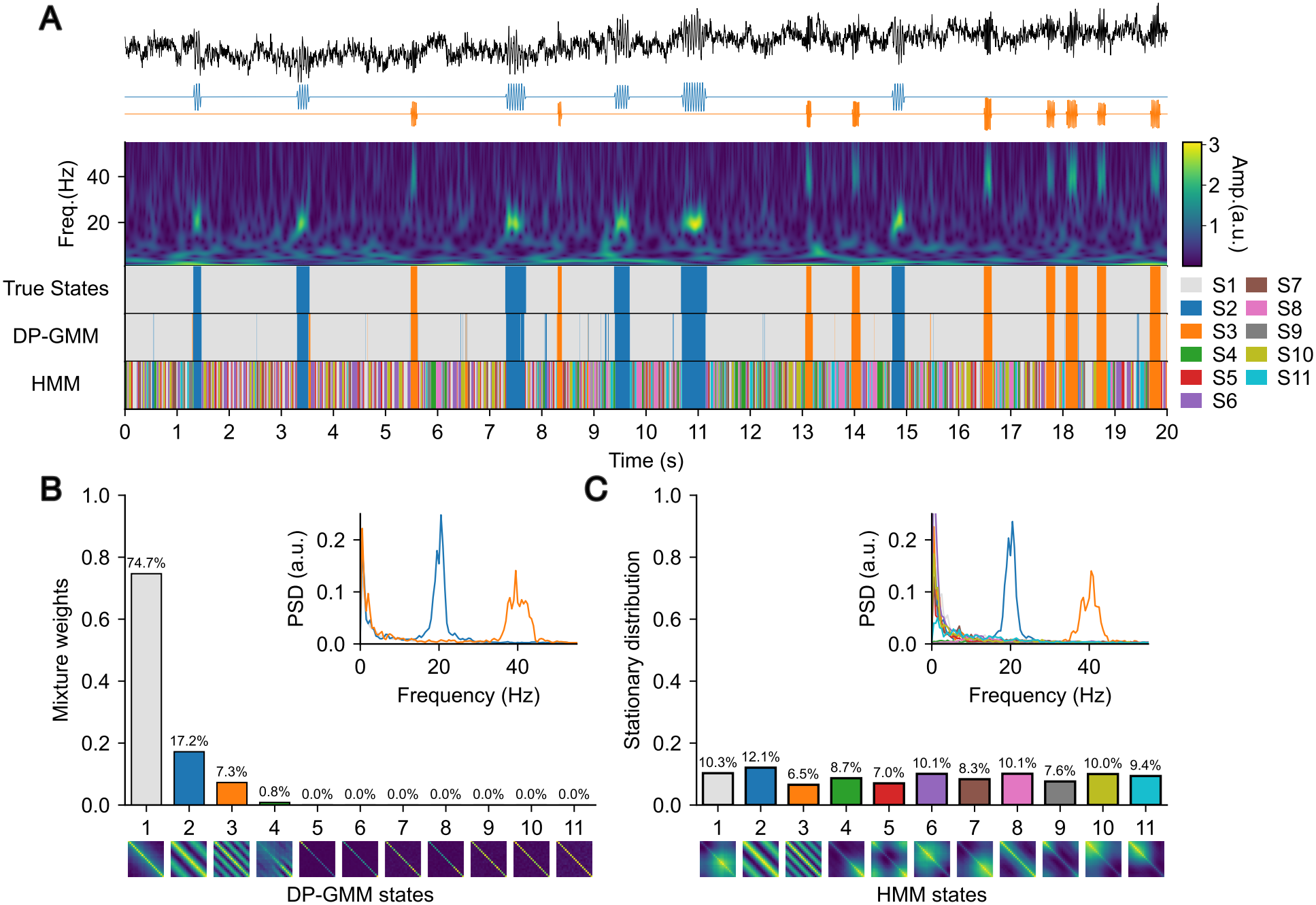
The DP-GMM recovered the true number of states and achieved high clustering accuracy by eliminating inactive components. (A) Representative snippet from a three-state synthetic signal containing transient 20 Hz (blue) and 40 Hz (orange) oscillatory states at SNR = 2 dB. The panel shows the panel shows the DP–GMM and HMM state sequences before imputation. (B) Outputs of the DP-GMM: posterior mixture weights and autocovariance matrices, with state-wise power spectra computed for the active states. (C) Corresponding outputs for the HMM, including the stationary distribution. We computed the stationary distribution ***π***_*∞*_ by solving 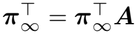 subject to 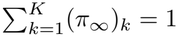, where ***A*** is the estimated transition matrix. Note that ***π***_*∞*_ does not have the same interpretation as the DP-GMM mixture weights: it reflects the long-run fraction of time the Markov chain spends in each state under ***A***, rather than the probability of a component generating an observation independently of temporal dynamics.

Although some of the HMM’s autocovariance matrices exhibited nonuniform diagonal values, suggesting that the corresponding states were edge-like, these states did not appear to occur exclusively at the boundaries of state segments, unlike the DP-GMM’s edge states (Fig. S2). Moreover, model selection based on the lowest NELBO did not identify three states as optimal for the HMM (Fig. S10). Instead, the NELBO continued to decrease as the number of states increased throughout the tested range, up to 21 states. No clear elbow was apparent, and even candidate elbows around 8 or 12 states would still overestimate the true number of states.

This comparison demonstrates the practical benefit of adapting the number of active states when the true state count is unknown. By assigning negligible mixture weights to redundant states, the DP-GMM reduces the risk of inaccurate state estimation when practitioners specify more states than are present in the data.

### The DP-GMM is robust to moderate amplitude differences and waveform asymmetry

Having established that the DP-GMM can identify frequency-selective states in moderately noisy signals, we next examined its sensitivity to variations in oscillation amplitude and morphology.

We generated 20 independent three-minute simulations containing two states: a 20 Hz oscillatory state and a noise state. Signals were sampled at *f*_*s*_ = 250 Hz. We used 33 lags, selected to approximate a target embedding window of 0.125 s, corresponding to 2.5 cycles at 20 Hz.

### The DP-GMM is tolerant of amplitude differences

We first examined whether differences in burst amplitude were sufficient to produce separate states. We fixed the lower-amplitude regime at an SNR of 2 dB and systematically increased the amplitude of the higher-amplitude regime relative to this baseline. The amplitude ratio was increased in increments of 0.5, up to a maximum of 4.

At lower amplitude ratios, the model assigned low- and high-amplitude 20 Hz oscillations to a single oscillatory state (Fig. S11). However, it also identified many edge states at segment boundaries. This was consistent with our earlier observation that the model tends to identify edge states at sufficiently high SNRs (Fig. S4), as higher oscillation amplitudes corresponded to higher SNRs in these simulations. After post-hoc removal of edge states, only two states remained for amplitude ratios up to 2.5. Separation between low- and high-amplitude oscillations began to emerge at an amplitude ratio of 3. At ratios above 3.5, three states remained after removing edge states, corresponding to high-amplitude beta oscillations, low-amplitude beta oscillations, and noise.

These results indicate that the DP-GMM can distinguish oscillatory events with sufficiently large amplitude differences, while grouping oscillations with moderate amplitude variations into a common oscillatory state. This can be advantageous when the primary goal is to identify oscillatory activity based on its spectral characteristics rather than differences in amplitude. Whether amplitude variations should be represented by separate states depends on the research question and the properties of the data. When amplitude variability is of particular interest, the identified oscillatory states can serve as a basis for subsequent analyses of within-state amplitude dynamics.

### The DP-GMM is stable under non-sinusoidal signals

Neural oscillations can exhibit non-sinusoidal waveforms with unequal rise and decay durations. To evaluate how this asymmetry affects DP-GMM clustering performance, we compared pure sinusoids with asymmetric sine waves.

We generated the asymmetric sine waves using the NeuroDSP function `sim_oscillation` with `cycle=‘asine’` [55]. The rise–decay symmetry parameter (`rdsym`) specifies the fraction of each cycle spent in the rising phase. A value of 0.5 produces a symmetric sine wave. At 0.2, the waveform rises over 20% of the cycle and decays over the remaining 80%, giving it a sawtooth-like shape (Fig. S12A). For the clustering comparison, we sampled `rdsym` uniformly between 0.1 and 0.9 for each oscillatory segment. Clustering performance and estimated state counts were comparable between signals with asymmetric and sinusoidal waveforms, suggesting that the DP-GMM was robust to the tested waveform variations (Fig. S13). Although waveform asymmetry varied across oscillatory segments within each signal, the model did not separate these segments into shape-specific subclasses. Occasionally, the model identified additional states that were different from edge states but also had covariance matrices with nonuniform diagonal values (e.g., state S3 in Fig. S13B). The default CV criterion removed these additional states.

Overall, the DP-GMM accurately distinguished an oscillatory state from a noise state even under variable waveforms. Within the range tested, differences in rise–decay asymmetry did not lead the model to infer separate substates, likely because the different waveforms still had similar autocorrelation structures.

### The DP-GMM separates states with different aperiodic components

We next shifted our focus from finding oscillations to non-oscillatory aperiodic backgrounds, treating what we had previously termed “noise” as a part of signal components worth characterizing. We hypothesized that differences in aperiodic structure could be captured by the DP-GMM’s state-specific covariance matrices, allowing the model to distinguish states with different aperiodic exponents.

For each exponent condition, we generated 20 five-minute simulations sampled at *f*_*s*_ = 250 Hz and an SNR of 2 dB. Each simulation contained four ground-truth states: an aperiodic state with exponent *β*_1_ = 1 (pink noise); a 20 Hz oscillatory state superimposed on this background; a second aperiodic state with exponent *β*_2_; and a 20 Hz oscillatory state superimposed on the second background (Fig. S14A). We varied *β*_2_ from 0 to 2 in increments of 0.2, excluding *β*_2_ = 1, giving ten exponent conditions. The power spectral densities, averaged across the 20 simulations for each condition, showed progressively steeper aperiodic slopes with increasing *β*_2_ (Fig. S14B).

Within each five-minute simulation, the two aperiodic backgrounds alternated in 30-s blocks, with five repetitions of each block pair. Each block contained oscillatory bursts interspersed with periods containing only the aperiodic background. For each block, the background amplitude was scaled to achieve a target SNR of 2 dB, calculated using the mean burst power during oscillatory periods and the variance of the background after 0.5 Hz high-pass filtering. The scaled, unfiltered background was then added to the bursts.

In the four-state simulation, the DP-GMM achieved excellent clustering accuracy (ARI = 0.90 before imputation; ARI = 0.92 after imputation; Fig. 5A), while retaining four active states from the initial set of candidate states (Fig. 5B). The model accurately distinguished aperiodic components with exponents *β*_2_ *<* 0.4 or *β*_2_ *>* 1.2 from the baseline component with *β*_1_ = 1. Even at *β*_2_ = 0.8 and *β*_2_ = 1.2, which were close to the baseline exponent, the DP-GMM maintained separation between the two aperiodic components while identifying a single 20 Hz oscillatory state.

**Fig 5.**
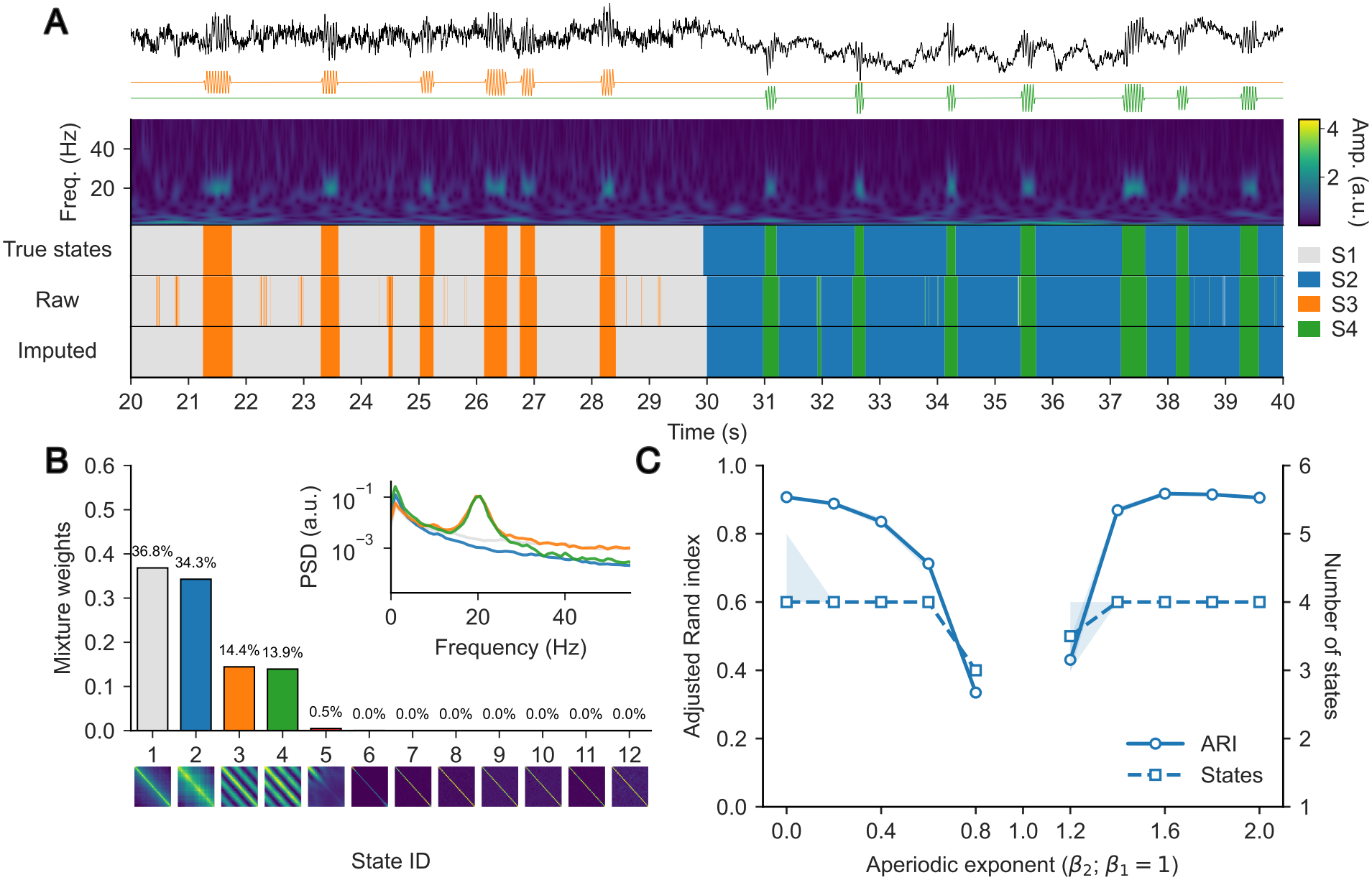
The DP-GMM distinguished aperiodic states with different exponents. (A) An example simulated signal containing aperiodic states with exponents *β*_1_ = 1.0 (grey) and *β*_2_ = 1.6 (blue), and 20 Hz oscillatory states embedded in aperiodic backgrounds with exponents *β*_1_ = 1.0 (orange) and *β*_2_ = 1.6 (green) at an SNR of 2 dB. The raw decoded state sequences are shown alongside those obtained after post-hoc *k*-NN imputation. (B) Posterior mixture weights alongside the corresponding state autocovariance matrices.(C) Adjusted Rand index and number of estimated states as a function of *β*_2_, with *β*_1_ fixed at 1.0. We excluded the benchmark condition *β*_2_ = *β*_1_ = 1.0 because it reduces to a two-state scenario consisting of one 20 Hz oscillatory state and one aperiodic state.

These simulations demonstrate that the model can not only distinguish between aperiodic states but also separate oscillatory states that share the same frequency but differ in their underlying aperiodic structure. The DP-GMM was therefore sensitive to both periodic and aperiodic signal characteristics, and state separation extended beyond differences in oscillatory frequency.

### The DP-GMM revealed multiple transient oscillatory and aperiodic states in resting MEG

Having validated the time-delay embedded DP-GMM on synthetic data with known ground truth, we next applied it to a single parcel in the left motor cortex of MEG recordings in a publicly available dataset [57]. To determine the embedding dimension, we first obtained the candidate frequency components. Here, we identified five candidate frequencies: 5, 11, 17, 21, and 32 Hz based on the peaks of the group-averaged power spectral density (PSD) (Fig. S15). Importantly, these candidate peaks were used only as a pragmatic guide for choosing the embedding window and *k*-NN imputation window. Here, we used 15 embeddings derived from 2.5 cycles of the average frequency of 17 Hz, covering a 0.144 s window. We then used the DP-GMM with 15 states (⌈ln 1929818⌉). As in the synthetic-signal benchmark, extremely brief state segments were common in the estimated state sequences and dominated the lifetime distributions, with median state lifetimes of 10–20 ms across all states (Fig. S16). To reduce this fragmentation, we applied *k*-NN imputation with a 30 ms window.

The DP-GMM decomposed resting motor-cortex activity into two aperiodic states (States 1 and 2) and five short-lived oscillatory states (States 3–7). The inferred state sequence was largely occupied by one aperiodic state (State 1), intermittently interrupted by brief activations of both another aperiodic state (State 2) and the oscillatory states that aligned with localized increases in the wavelet representation (Fig 6A). These seven active states accounted for 98.77% of the total posterior mixture weight, while the remaining components had negligible weights except for one state, which was additionally treated as inactive due to its high CV of the covariance diagonal. The posterior mixture weights for the seven active states were then renormalized to sum to 1 (Fig 6B). The estimated covariances for the oscillatory states (States 3–7) exhibited characteristic striped patterns consistent with oscillatory structure. They exhibited a narrowband peak at 12 Hz (State 3), 26 Hz (State 4), 20 Hz (State 5), 8 Hz (State 6), and 22 Hz (State 7) (Fig 6C). These oscillatory states aligned broadly with the prominent PSD candidates (Fig. S15). Although some oscillatory states had relatively similar peak frequencies (e.g., States 3 and 6 at 12 and 8 Hz, respectively, and States 5 and 7 at 20 and 22 Hz, respectively), their corresponding PSDs differed mainly in magnitude. Consistent with these observations, the state-wise autocovariance functions (ACFs), derived from the estimated autocovariance matrices, also differed in magnitude (Fig. S17). In contrast, State 1 showed no clear narrowband peak, whereas State 2 exhibited a pronounced 1*/f* -like spectrum with maximal power at 2.5 Hz. States 1 and 2 also showed no clear oscillatory structure in their ACFs (Fig. S17).

**Fig 6.**
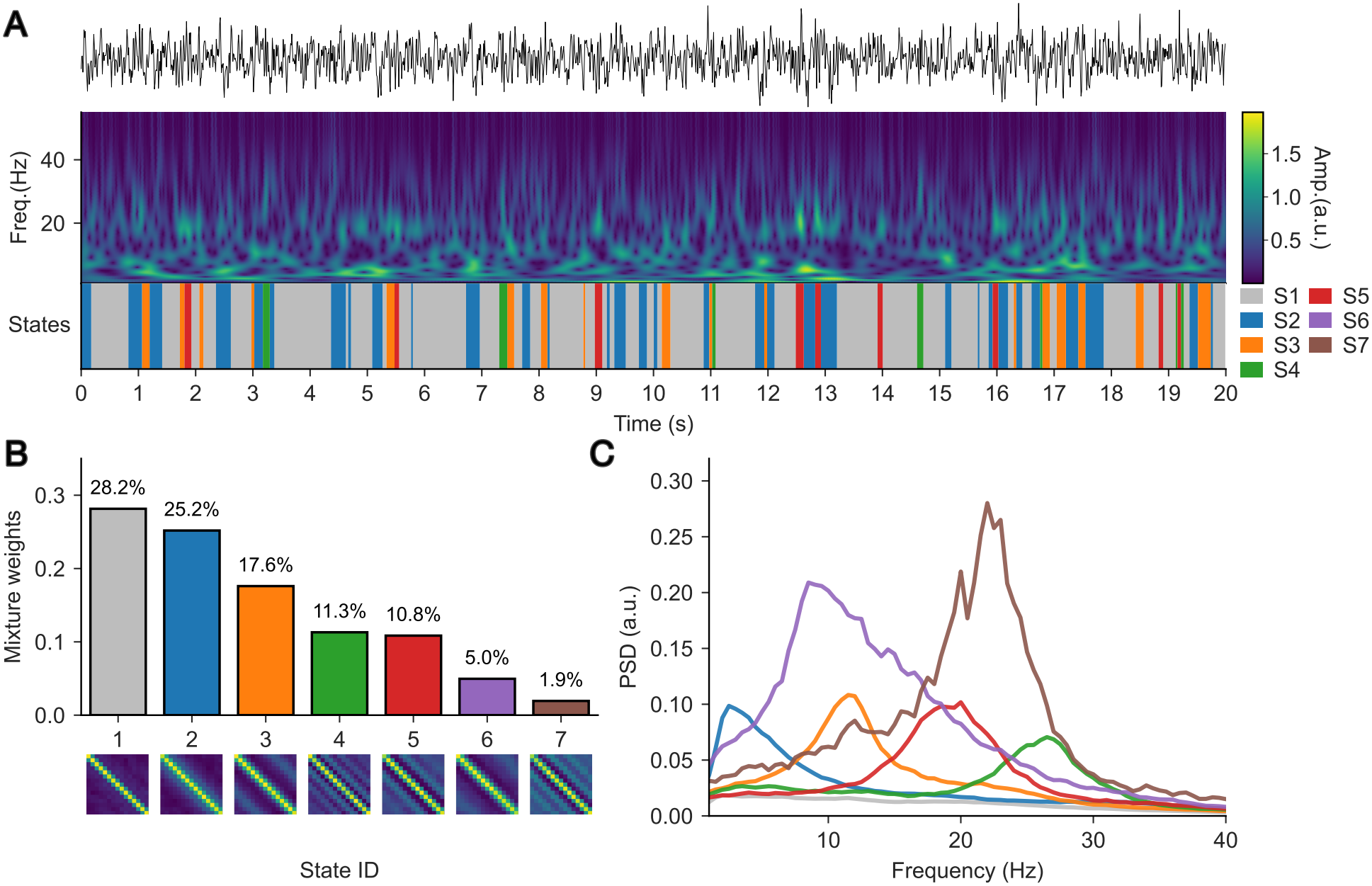
The DP-GMM identified seven spectrally distinguishable neural states in resting motor-cortex MEG activity. (A) Decoded state sequence for the seven active states after post-hoc *k*-NN imputation. (B) Posterior mixture weights for the active components, shown alongside the corresponding state autocovariance matrices. (C) State-wise power spectra computed from the seven active states.

Having identified a common set of oscillatory and aperiodic states at the group level, we next examined how these states were expressed across individual participants. Fractional occupancy varied substantially across participants for all states (Fig 7A). Among participants exhibiting each state, median fractional occupancy [Q1, Q3] was: State 1, 38.42% [25.11%, 52.52%]; State 2, 24.80% [15.40%, 34.96%]; State 3, 12.62% [9.12%, 18.33%]; State 4, 6.75% [2.15%, 11.07%]; State 5, 4.81% [2.90%, 10.55%]; State 6, 3.29% [1.53%, 5.28%]; and State 7, 0.54% [0.14%, 1.60%]. States 1–5 were present in all 65 participants, whereas States 6 and 7 were present in 64/65 (98.5%) and 56/65 (86.2%) participants, respectively. To further characterize the identified states, we also summarized state lifetimes, state power, occurrence rates, and CVs of inter-state intervals across participants (Fig 7B). The dominant baseline state (State 1) persisted longest, with a median lifetime [Q1, Q3] of 170 [130, 210] ms, whereas the other aperiodic state (State 2) had a shorter median lifetime of 120 [100, 140] ms. The oscillatory states exhibited shorter, burst-like episodes, with median lifetimes of 90 [80, 110], 85 [75, 100], 90 [70, 100], 105 [90, 130], and 70 [60, 96.25] ms for States 3–7, respectively. At their respective peak frequencies (11.5, 26.5, 20, 8.5, and 22 Hz), these lifetimes corresponded to approximately 1.03, 2.25, 1.80, 0.89, and 1.54 cycles. Notably, States 6–7 expressed comparatively high power (median power: State 6, 2.93 a.u.; State 7, 2.97 a.u.) despite their low occupancy. Occurrence rates broadly reflected the occupancy structure: States 1–3 occurred frequently, with median occurrence rates of 1.2–1.8 Hz, whereas States 4–7 occurred less frequently, with median occurrence rates of 0.06–0.64 Hz. However, occurrence rates showed substantial variability across participants. Median CVs of inter-state intervals ranged from 1.03 to 1.31, consistent with irregular state onsets.

**Fig 7.**
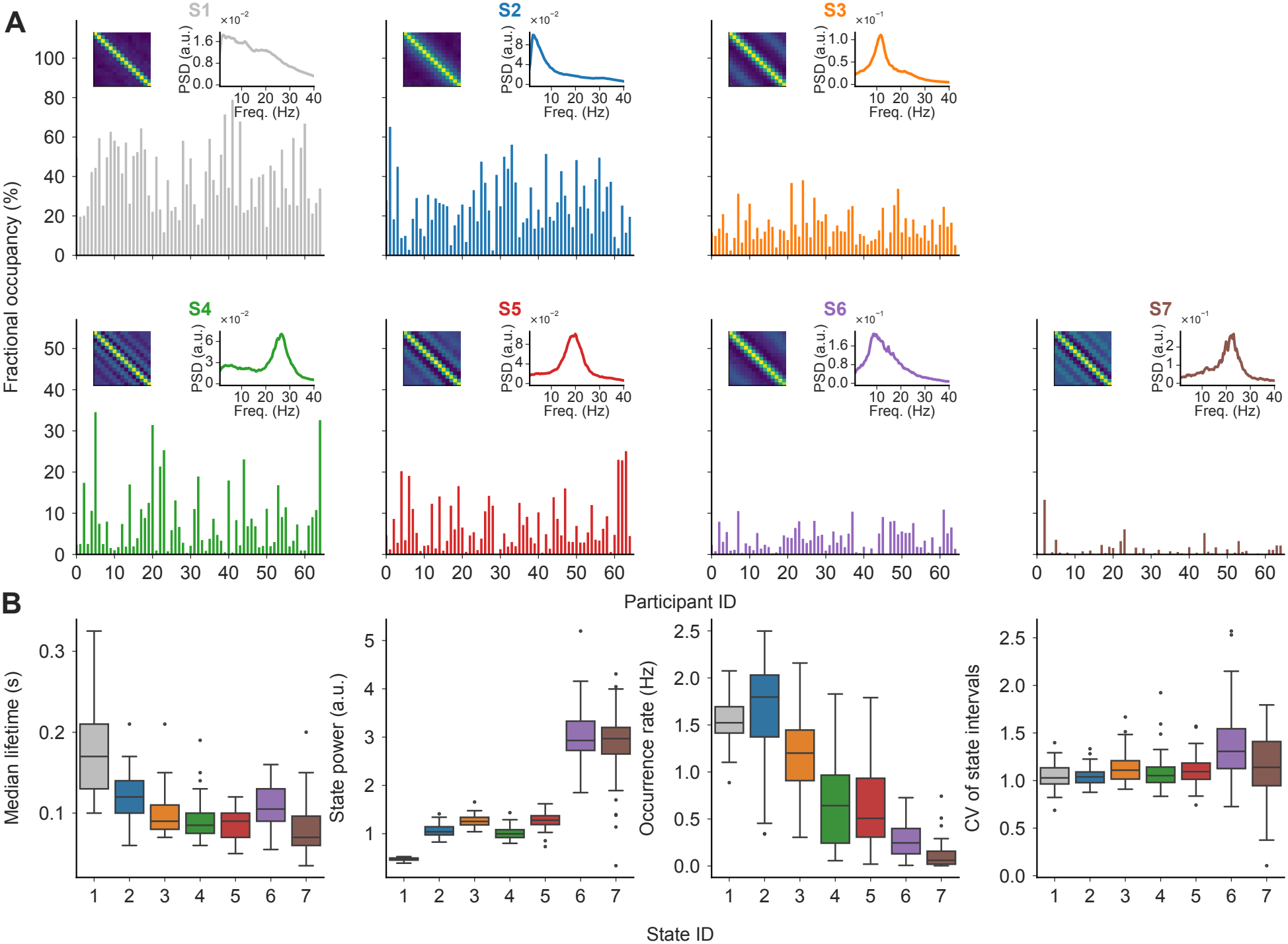
Participant-level summaries for the seven MEG states. (A) Fractional occupancy across participants (*N* = 65) for each of the seven active states. Insets show the state-specific autocovariance matrices and power spectra. (B) Boxplots of median lifetime, state power, occurrence rate, and coefficient of variation (CV) of inter-state intervals for each state. For the lifetime boxplots, we first calculated the median lifetime of each state within each participant. The boxplots summarize these participant-level medians, with the central line indicating the median across participants.

We examined whether the spectral characteristics of the inferred states changed with embedding dimension (*E*). The found states showed broadly similar spectral profiles and autocovariance patterns. However, their time courses were less strongly correlated at *E* = 11, 13, and 17, where the model retained eight states and separated one state into two substates (Fig. S18).

For comparison, we fitted a seven-state HMM to the same resting-state MEG data (Fig. S19). HMM states S1 and S5 had similar state spectra resembling those of the two aperiodic DP-GMM states. The HMM also identified comparable oscillatory states with alpha- and beta-band activity. Within both bands, however, the HMM state spectra appeared broader and overlapped more than those obtained with the DP-GMM, making the individual oscillatory states harder to distinguish (Fig. 6C and Fig. S19C). Finally, we examined whether the negative ELBO (NELBO) could guide model-order selection. As in the simulations, the NELBO decreased overall as *K* increased, despite several local increases, and reached its minimum at the largest model order tested (*K* = 21; Fig. S20).

Overall, the time-delay embedded DP-GMM revealed multiple transient oscillatory and aperiodic states in the resting-state MEG data. These states differed in spectral characteristics such as peak frequency, power, and aperiodic structure.

## Discussion

This work demonstrates that the time-delay embedded DP-GMM enables the unsupervised identification of multiple neural states with distinct spectral characteristics without specifying the number of states in advance. Its Dirichlet-process prior shrinks the weights of redundant components towards zero, reducing sensitivity to model-order misspecification. This model not only accurately detects oscillations across multiple frequency bands but also distinguishes between aperiodic components with different exponents. We also applied this method to a resting-state motor-cortex MEG dataset. In these data, the model identified several brief oscillatory states with lifetimes equivalent to only a few cycles, alongside transient aperiodic states with different spectral features.

The DP-GMM was particularly discriminative to differences in both oscillatory frequency and aperiodic structure, as shown by the synthetic and empirical benchmarks. It distinguished states with closely spaced oscillatory frequencies, such as 35 and 40 Hz, even in noisy signals (e.g., SNR = 0 dB; Fig. 3C). It also separated states whose aperiodic components differed only slightly in their exponents (e.g., *β*_1_ = 1 and *β*_2_ = 0.8 or 1.2)Fig. 5C. This ability to characterize aperiodic dynamics is relevant because a growing body of literature suggests that aperiodic activity carries physiologically meaningful information and is associated with cognitive function, neurodevelopmental stages, and clinical conditions [14, 58–61], rather than merely reflecting background noise.

The DP-GMM was less sensitive to variations in burst amplitude. It tended to assign oscillatory events at the same frequency to a single state unless their amplitudes differed substantially (Fig. S11). In the MEG dataset, states 5 and 7 had spectral peaks at 20 and 22 Hz, respectively (Fig. 6C) . Their separation likely reflected a combination of a large amplitude difference and subtler differences in oscillatory frequency and aperiodic structure. In contrast, bursts with different durations and morphologies (Fig. S13) were assigned to the same state under the tested conditions. Because the DP-GMM distinguishes states according to their autocovariance structure, these variations may not have altered that structure sufficiently to produce separate states.

This relative insensitivity may be advantageous because it can limit unnecessary state subdivisions and simplify interpretation. The DP-GMM can first identify states based primarily on oscillatory frequency and aperiodic structure, after which amplitude dynamics, state lifetimes, and burst morphology can be characterized in post-hoc analyses. This can be complemented by existing algorithms, such as specparam (formerly FOOOF) [59], CHO [62] and cycle-by-cycle analysis [63] to characterize oscillatory events and waveform morphology. State lifetimes can be quantified directly from the inferred state sequence.

Characterizing burst morphology is important because waveform shape can provide physiological information that complements spectral descriptions [64, 65]. For example, beta burst waveform diversity has been proposed to reflect characteristic patterns of synaptic input to cortical circuits [13].

Although the DP-GMM recovered the correct number of states across many simulation conditions, it tended to infer a single state at SNR = − 10 dB. This outcome was expected because the background noise largely masked the oscillations that distinguished the simulated states. We consider this behavior useful because the model favored a single noise-dominated state when the data provided little evidence for additional states. At high SNRs, the DP-GMM tended to overestimate the number of states by assigning additional states to time points near the boundaries of visits to the dominant states. These additional states typically had covariance matrices with nonuniform diagonal elements and could be excluded using post-hoc criteria based on the coefficient of variation of those elements. The number of inferred states also tended to increase with recording duration (Fig. S21). We attribute this increase primarily to the model’s ability to resolve subtler differences in longer recordings, including patterns at state boundaries that were otherwise apparent mainly at high SNRs.

We approximated the infinite DP mixture by truncating its stick-breaking representation at *K*_upper_ = ⌈ln *N* ⌉ components, where *N* is the number of signal samples. For three-minute recordings sampled at 250 Hz, this rule gave *K*_upper_ = 11. At a fixed reference concentration of *α* = 1, the theoretical bound *ϵ* on the *L*_1_ distance between the truncated and infinite mixture marginal densities was uninformative at this cap and first fell below 0.01 at *K*_upper_ = 25 [39]. In practice, increasing the cap from 11 to 27, the first tested value above this threshold, had little effect on clustering accuracy (Fig. S22). Because larger upper bounds can increase memory requirements when fitting long neural recordings, we still recommend using the log rule as a practical starting point. If all components remain active under the mixture-weight criterion before edge-state removal, we recommend increasing *K*_upper_ and refitting the model, so that the model can estimate how many states are active within this maximum.

We used stochastic variational inference to approximate the posterior distributions of the model parameters for both the DP-GMM and the HMM. The mean-field Gaussian approximation used here may underestimate uncertainty in the inferred parameters. Although we used only the posterior medians in subsequent analyses, examining the approximate posterior distributions could provide further insight into parameter uncertainty. For instance, the posterior distribution of the mixture weights may help us quantify uncertainty about the number of retained states. By contrast, when only point estimates are needed, maximum a posteriori estimation could simplify inference and reduce optimization time. Future studies should investigate these possibilities.

The thresholding method performed relatively consistent across frequency conditions and slightly better under very noisy conditions than the DP-GMM. This advantage was expected because thresholding was applied to narrow band-pass-filtered signals centered on the known simulated target frequencies. Such filtering increases the effective SNR within the target band [66], facilitating the detection of band-limited events even in very noisy signals. When the target frequency band is known in advance or can be independently justified, thresholding may suffice for burst detection without the added complexity of the DP-GMM. In practice, however, selecting and justifying target bands is nontrivial. Peak frequencies and bandwidths can vary across participants [67] and brain regions [68], making fixed detection bands difficult to justify. Even when a target band is justified, global amplitude thresholds can be poorly calibrated when oscillation amplitude varies substantially across time and recordings, leading to missed low-amplitude events or inflated false positives [22]. Multiple oscillatory components further complicate band selection and threshold calibration. The thresholding method may miss relevant components or misclassify broadband aperiodic activity as oscillatory activity [14]. Moreover, the thresholding method used here focuses on detecting oscillatory events within predefined detection bands and does not explicitly characterize aperiodic activity.

In the synthetic benchmarks, both the DP–GMM and the HMM occasionally assigned brief segments to the wrong state, possibly because the signal pattern within an embedding window resembled that of another state. The DP–GMM tended to produce many brief state fragments that would dominate state lifetime estimates without post-hoc *k*-NN imputation (Fig. S16). This is mainly because the DP-GMM assigns states independently across time given the mixture parameters. The HMM, on the other hand, exhibits fewer but longer fragments. This is most likely because the HMM enforces temporal regularization through a transition matrix, which can discourage rapid state switching [23], but can also prolong erroneous state activations once they occur. These slightly longer spurious assignments can similarly underestimate state lifetime. Although increasing the *k*-NN window could further suppress false positives for the HMM, it would also raise the risk of reassigning true oscillatory events to neighboring states, thereby increasing false negatives. Using posterior state probabilities in place of hard state assignments may reduce the influence of brief state visits. Hard classification assigns each time point to a single state, even when that state is only slightly more probable than the alternatives. Posterior weighting accounts for this uncertainty by allowing each time point to contribute to multiple states in proportion to their probabilities.

A complementary approach is to model state durations explicitly. Hidden semi-Markov models (HSMMs) allow each state to have its own duration distribution, relaxing the geometric duration assumption of standard HMMs [69]. Assigning low probability to unrealistically short durations may reduce the number of spurious brief visits in the inferred state sequence. Bayesian nonparametric extensions include the hierarchical Dirichlet process HMM (HDP-HMM) [70] and its semi-Markov counterpart, the HDP-HSMM [71]. Like the DP-GMM, both models use Dirichlet process priors that allow the number of occupied states to adapt to the data. The HDP-HSMM combines this flexibility with explicit modeling of state persistence. Future work should compare the DP-GMM and the HDP-HSMM on the same time-delay embedded data to assess whether the HDP-HSMM yields fewer spurious brief visits.

We observed that the choice of embedding dimension (i.e., the number of lags) strongly influences clustering performance for both the DP-GMM and the HMM. As a pragmatic heuristic, we set the embedding window to span approximately 2.5 cycles at the frequency of interest for the DP-GMM and the HMM. This simple cycle-based rule can break down at very slow or very fast frequencies, where it can yield an impractically large or small number of lags. Moreover, in multi-frequency settings, any single embedding choice is necessarily a compromise. This compromise can be suboptimal when a recording contains both low- and high-frequency oscillations. This may help explain why both models showed lower clustering accuracy for the 10/40-Hz pair than for pairs with intermediate frequency separations (Fig. S7). The selected window may be too short to capture enough cycles of the 10-Hz oscillation, yet long enough to blur the boundaries of brief 40-Hz events. This reflects a trade-off between representing oscillatory structure and preserving temporal precision. Additionally, the choice of embedding dimensions still requires a rough estimate of the frequencies of interest, although nearby embedding dimensions yielded similar results in our analyses (Fig. S1 and Fig. S18). Several methods have been proposed for selecting embedding dimensions in state-space reconstruction (for a review, see [72]), but their applicability to oscillatory-state clustering remains an open question.

Our benchmarking simulations had several limitations. First, the studied range of oscillation frequencies were limited. Higher-frequency activity such as hippocampal ripple-range oscillations (∼ 80–250 Hz [73]) and high gamma (70–150 Hz [74]), as well as very low frequencies including delta (0.5–4 Hz) and theta (4–8 Hz [75]), remain to be explored. Second, we did not simulate nested or concurrent oscillations (e.g., high-frequency ripples nested within 10–16 Hz spindles during cortical slow oscillations, ∼ 0.5–1 Hz) [76]. Relatedly, our models assume a mutually exclusive state representation, in which each time point is assigned to a single latent state. By contrast, a recent approach, Dynamic Network Modes (DyNeMo), models activity as mixtures of modes, allowing overlapping mode contributions [24]. Such a model is a promising candidate for capturing nested or concurrent activity and may complement our Bayesian nonparametric approach. Third, our waveform simulations were limited to sinusoids and sawtooth-like oscillations. Future work should evaluate the DP-GMM using a broader range of physiological waveform shapes, including the arch-shaped sensorimotor mu rhythm [14, 64], and neocortical slow oscillations containing complex wave forms [64, 77]. Fourth, our assessment of the DP-GMM’s sensitivity to aperiodic structure was limited to variations in the spectral exponent. We did not systematically examine changes in other aperiodic parameters, such as the offset and knee frequency [59, 60]. Despite these simplifications, we expect the model to characterize a broader range of signal features because it exploits the signal’s autocovariance structure.

## Conclusion

Taken together, these results indicate that the time-delay embedded DP-GMM provides a robust, unsupervised framework for characterizing both oscillatory and aperiodic dynamics in noisy neural recordings, reducing reliance on narrow band-pass filtering and a priori specification of the number of states. The framework is therefore well suited for analyses in which the relevant signal components and the number of underlying neural states are unknown.

Conceptually, this approach redefines oscillation analysis from detecting activity within predefined frequency bands to discovering a sequence of multiple discrete neural states that differ in their spectral properties. This state-based perspective may facilitate more reproducible analyses of time-varying neural dynamics across participants, brain regions, and task contexts, and support investigations of how these dynamics relate to cognitive processes.

## Data and code availability

The code used for simulation, model fitting, and analysis is available in the TDE-DP-GMM GitHub repository. The simulated data (`simulations.zip`) and model benchmark results (`performance.zip`) are publicly available on Zenodo (DOI: 10.5281/zenodo.24430502).

We analyzed resting-state MEG data from the MEGUK partnership’s “CTF rest” dataset, recorded at the University of Nottingham using a 275-channel CTF system. Raw MEG recordings are available through the [MEGUK database](https://meguk.ac.uk/database/), subject to its data-access process. We do not redistribute raw MEG recordings.

We obtained the preprocessed, source-reconstructed MEG time series through the `osl-dynamics` workflow [24]. For each participant, we analyzed the time series from a single parcel in the left motor cortex. These time series are in the `bursts` folder of the `notts_meguk_giles.zip` archive on OSF. The accompanying GitHub repository provides the [preprocessing scripts](https://github.com/OHBA-analysis/osl-dynamics/tree/main/examples/toolbox_paper/ctf_rest/preproc) and the [archive download script](https://github.com/OHBA-analysis/osl-dynamics/tree/main/examples/toolbox_paper/ctf_rest/preproc).

## Acknowledgments

Parts of the computations were performed on the HPC cluster *Elysium* at Ruhr University Bochum, subsidized by the DFG (INST 213/1055-1).

## Supporting information

**Fig. S1.**
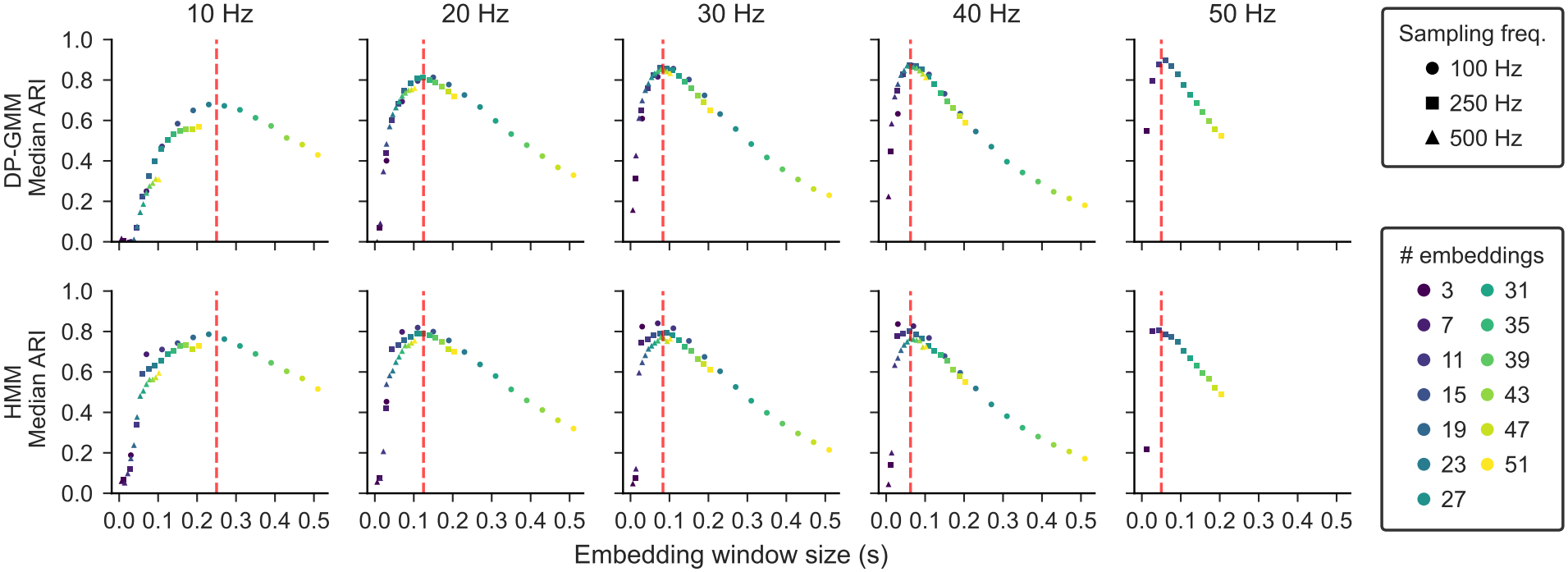
DP-GMM clustering performance peaked with an embedding window spanning approximately 2.5 cycles at the target frequency. We summarize the joint effect of *E* and *f*_*s*_ using the embedding-window duration *w*_*E*_ ≈ *E/f*_*s*_. Median ARI across five independent simulations is shown as a function of *w*_*E*_ for the DP-GMM (top row) and the HMM (bottom row) across target frequencies. The embedding dimension *E* is color-coded, and marker shape indicates the sampling rate. The vertical red dashed line indicates an embedding window spanning 2.5 cycles at the target frequency. Here, we simulated five independent 3-min samples per condition at SNR = +2 dB. For both the DP-GMM and the HMM, we used *K* = 2 states, assuming that the true number of states was known. Accordingly, the DP-GMM was not allowed to discard inactive states.

**Fig. S2.**
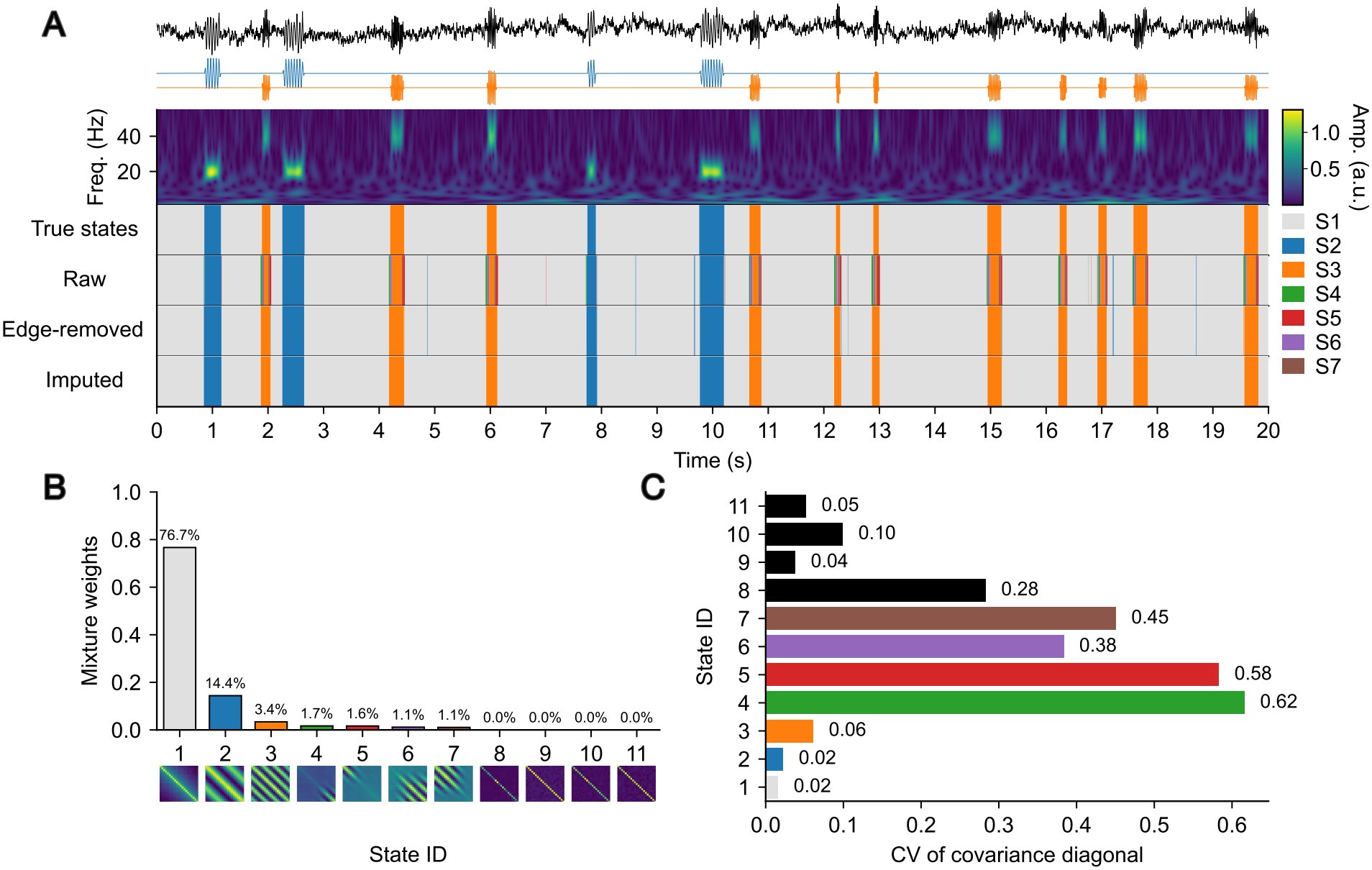
DP-GMM identifies edge states between noise and oscillatory periods. (A) Example raw trace from the three-state synthetic signal containing 20 Hz (blue) and 40 Hz (orange) oscillations at SNR = 6 dB, together with the wavelet amplitude (a.u.) and ground-truth state sequence. The raw estimated DP-GMM state sequence contained 7 active states out of 11 initial states after retaining 99% of the mixture weight. The edge-removed state sequence additionally removes excessive transitional states and retains three states, matching the true number of states. We then applied *k*-NN imputation to the edge-removed states. (B) Posterior mixture weights and autocovariance matrices. The autocovariance matrices for edge-related states typically show asymmetric diagonal patterns (e.g., S4–S11), whereas the desired states often have stable diagonal values (e.g., S1–S3). (C) Coefficient of variation (CV) of the covariance diagonal across all 11 states. The CV computes the normalized variability of covariance diagonal entries, with higher values suggesting greater asymmetry across lags. Here, edge states are defined as states with CV *>* 0.1 and are removed in addition to states with negligible weights.

**Fig. S3.**
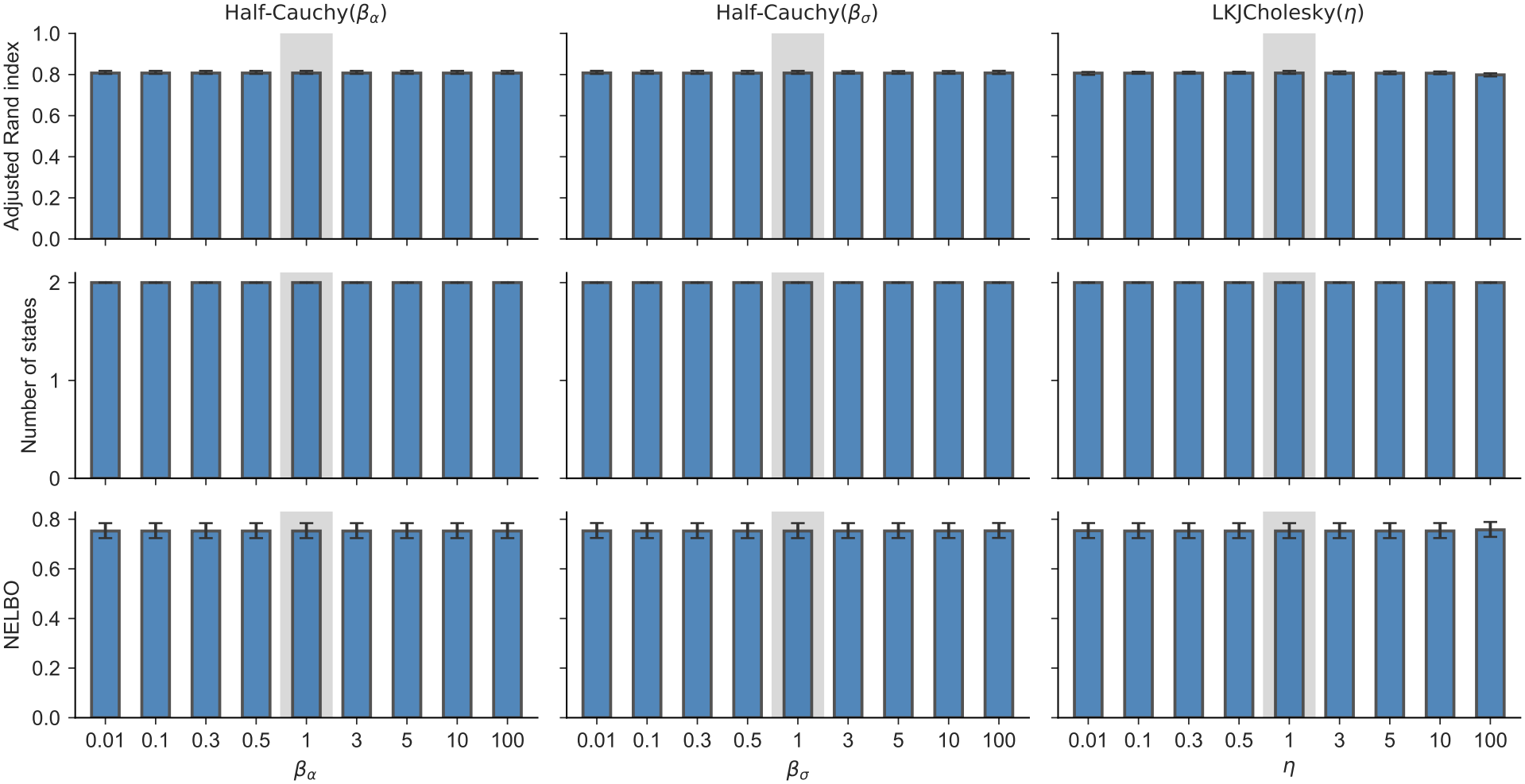
Sensitivity of the DP-GMM to prior hyperparameters. Columns show variation in the Half-Cauchy scale for the concentration parameter, *β*_*α*_ (left), the Half-Cauchy scale for the marginal standard deviations, *β*_*σ*_ (middle), and the LKJ correlation-prior concentration, *η* (right). Each parameter was varied individually while the other two were fixed at 1. Rows show the adjusted Rand index (top), number of active states (middle), and negative evidence lower bound (NELBO; bottom). Bars indicate medians across 20 independent simulations, and error bars span the interquartile range. Each simulation contained a 20 Hz oscillatory state and a pink-noise state at an SNR of 2 dB. Results are shown after mixture-weight truncation and before CV-based edge-state removal or *k*-NN imputation. Gray shading marks the default value of 1.

**Fig. S4.**
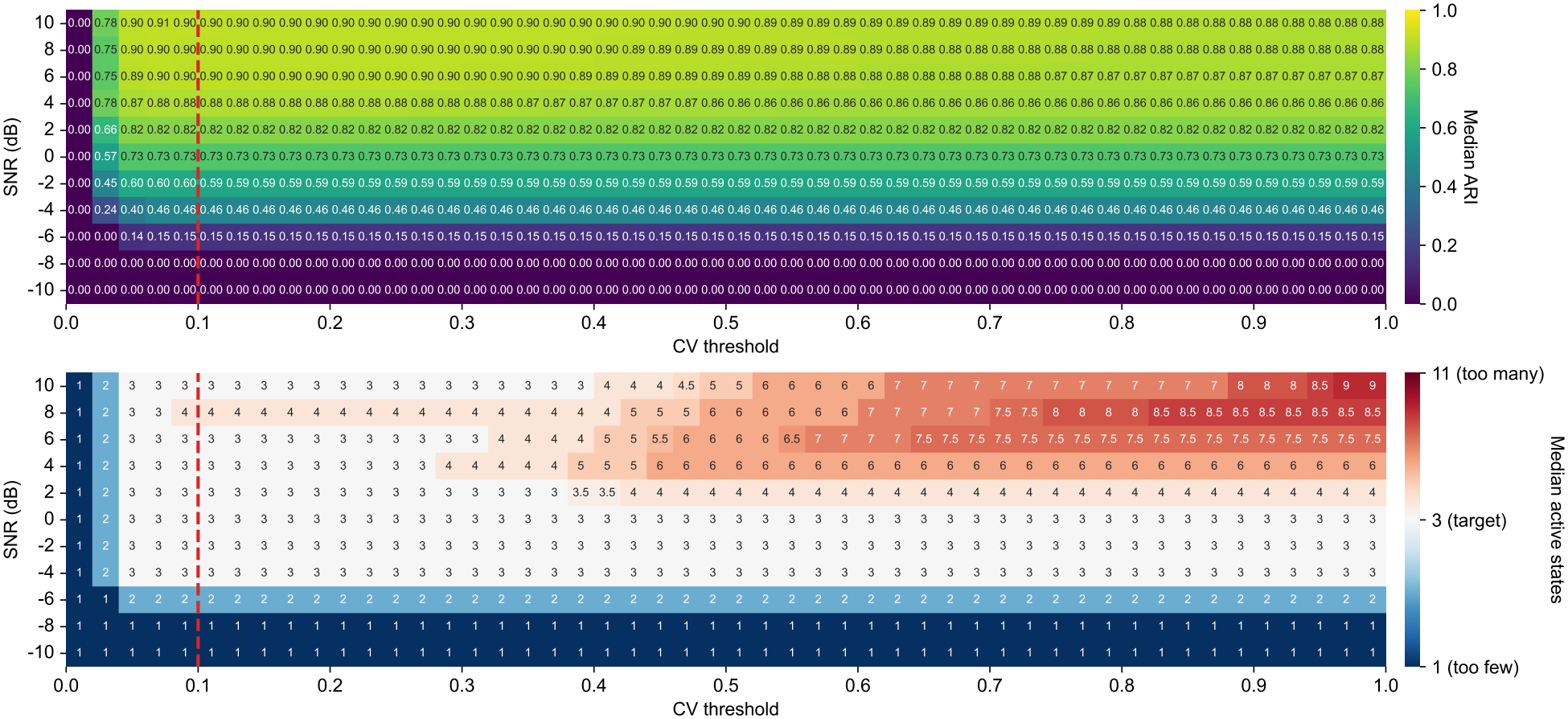
Sensitivity of DP-GMM clustering and state-count estimation to the CV threshold for edge-state removal. Heatmaps show the median adjusted Rand index (top) and median number of active states in the decoded sequence (bottom) for the three-state simulation containing 20 Hz and 40 Hz oscillations and a pink-noise state. Columns vary the CV threshold from 0.01 to 0.99 in increments of 0.02, and rows vary SNR from − 10 to 10 dB. For each threshold, components were first retained according to the 99% mixture-weight criterion, and components with a CV of covariance-diagonal entries above the threshold were then removed. State sequences were decoded again after this removal, without *k*-NN imputation. Cell annotations show the median values across simulations. The lower color scale is centered on the true number of states, *K* = 3, with blue indicating fewer states and red indicating more states. Red dashed lines mark the default CV threshold of 0.1.

**Fig. S5.**
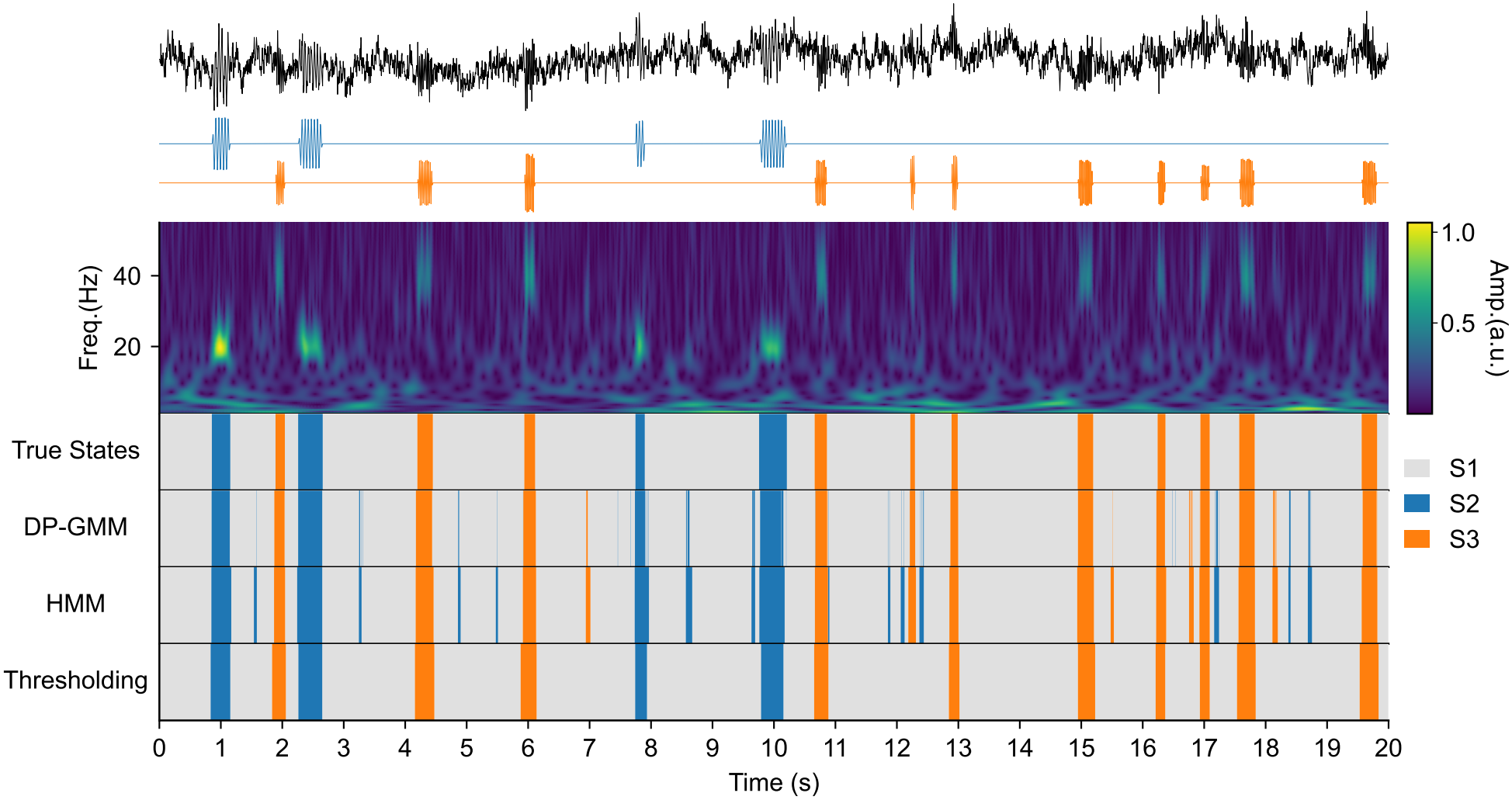
Brief state fragments are visible in the TDE-based model outputs. Rows show the three-state raw trace at SNR = 2 dB with 20 Hz (blue) and 40 Hz (orange) oscillations, wavelet amplitude, ground-truth states, and estimated states from DP-GMM, HMM, and the thresholding method *before k*-NN imputation.

**Fig. S6.**
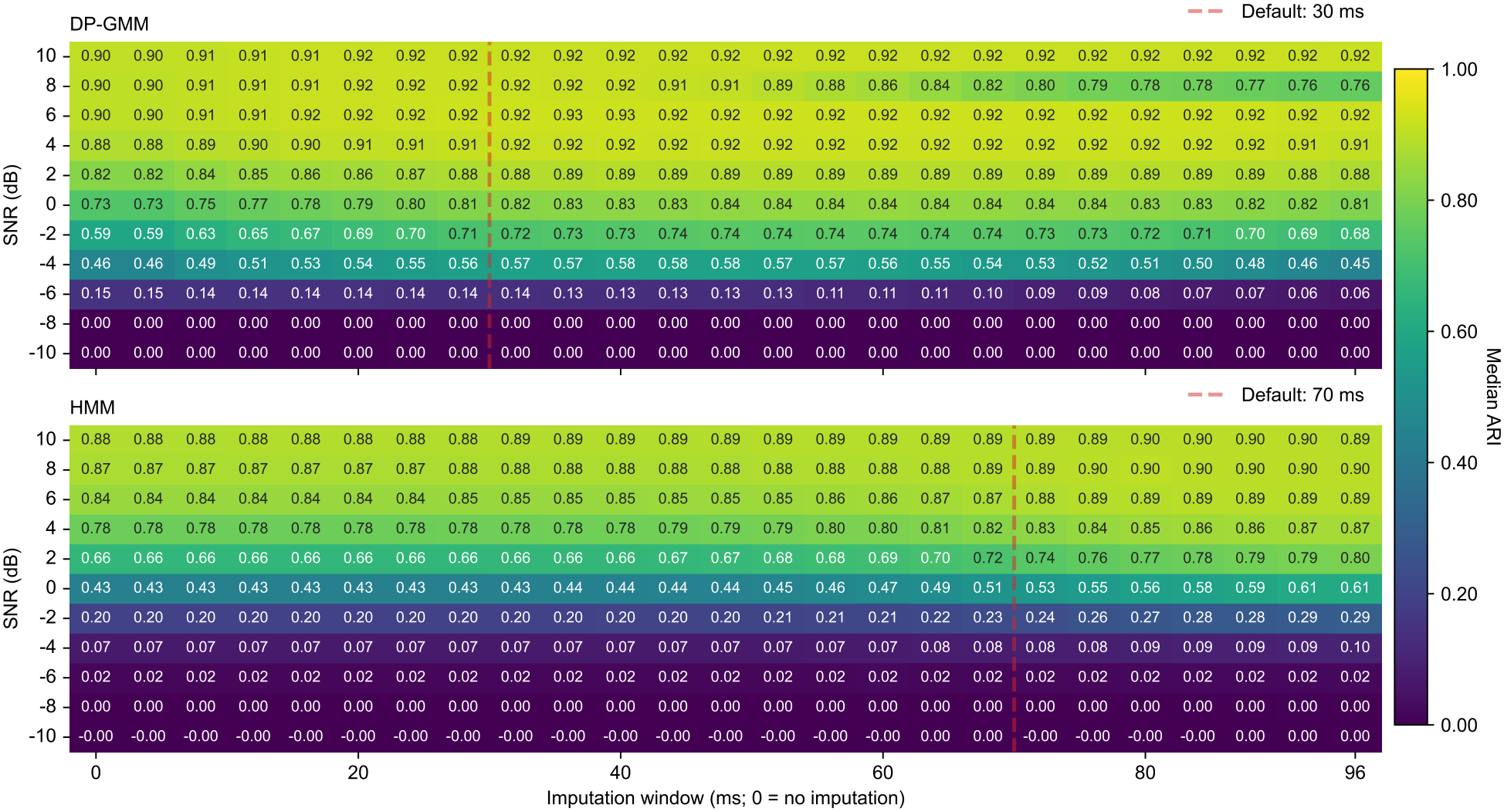
Sensitivity of DP-GMM and HMM clustering to the *k*-NN imputation window. Heatmaps show the median adjusted Rand index across 20 independent simulations per SNR for the DP-GMM (top) and HMM (bottom), using three-state signals containing 20 Hz and 40 Hz oscillations and a pink-noise state sampled at 250 Hz. DP-GMM state sequences were evaluated after CV-based edge-state removal. The imputation threshold was varied from 0 to 24 samples, corresponding to 0–96 ms in 4 ms steps; 0 denotes no imputation. For each threshold, shorter contiguous state segments were reassigned using *k*-NN imputation, starting from the same original decoded sequence. Cell annotations show median ARI, with a shared color scale for both methods. Red dashed lines mark the selected nominal defaults of 30 ms for the DP-GMM and 70 ms for the HMM.

**Fig. S7.**
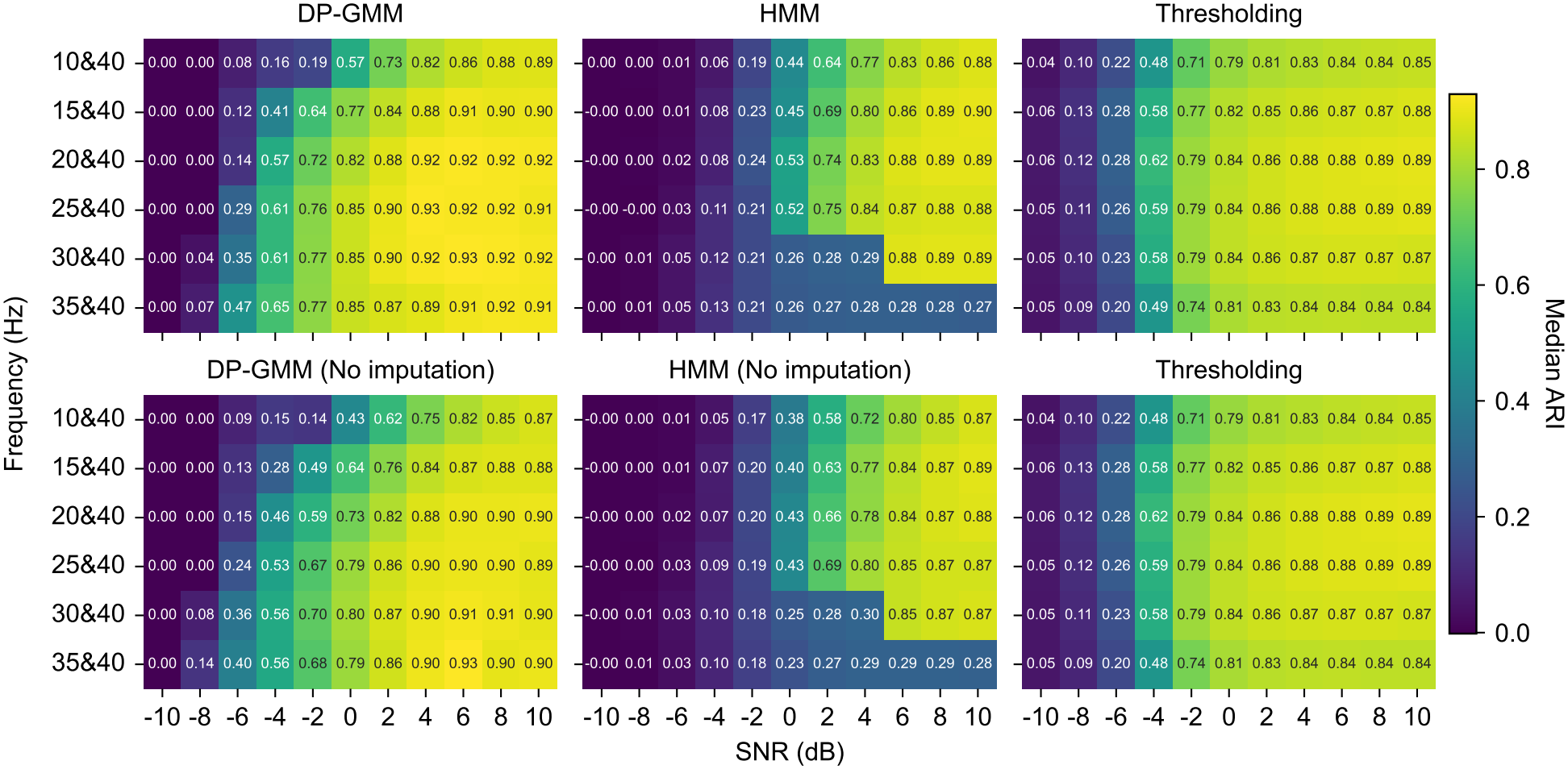
Performance heatmaps across SNR and frequency pairs with and without *k*-NN imputation. Median ARI heatmaps for the three-state scenario as a function of SNR and frequency conditions. Columns compare methods (DP-GMM, HMM, and thresholding). The top row shows results after post-hoc *k*-NN imputation of brief fragments, whereas the bottom row shows the corresponding results without imputation.

**Fig. S8.**
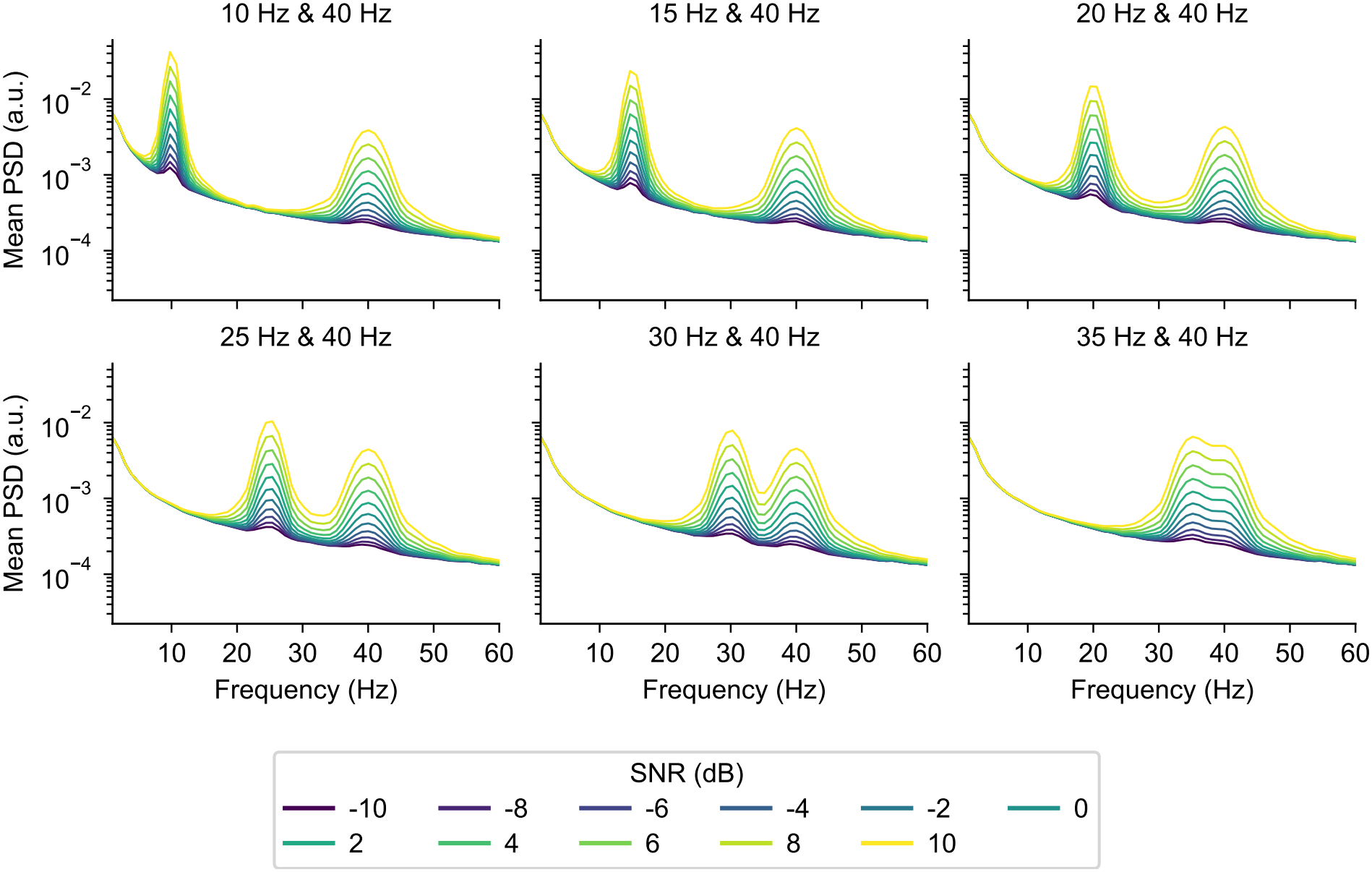
Power spectra of the three-state simulations across frequency pairs and signal-to-noise ratios. Mean power spectral densities (PSDs) of signals containing two oscillatory states and one pink-noise state. Panels show the six oscillatory frequency pairs: − 10 and 40, 15 and 40, 20 and 40, 25 and 40, 30 and 40, and 35 and 40 Hz. Colors indicate SNRs from 10 to 10 dB in 2-dB increments. PSDs were estimated separately for each signal using Welch’s method and averaged across 20 simulations per condition. Signals were sampled at 250 Hz, and power is shown on a logarithmic scale.

**Fig. S9.**
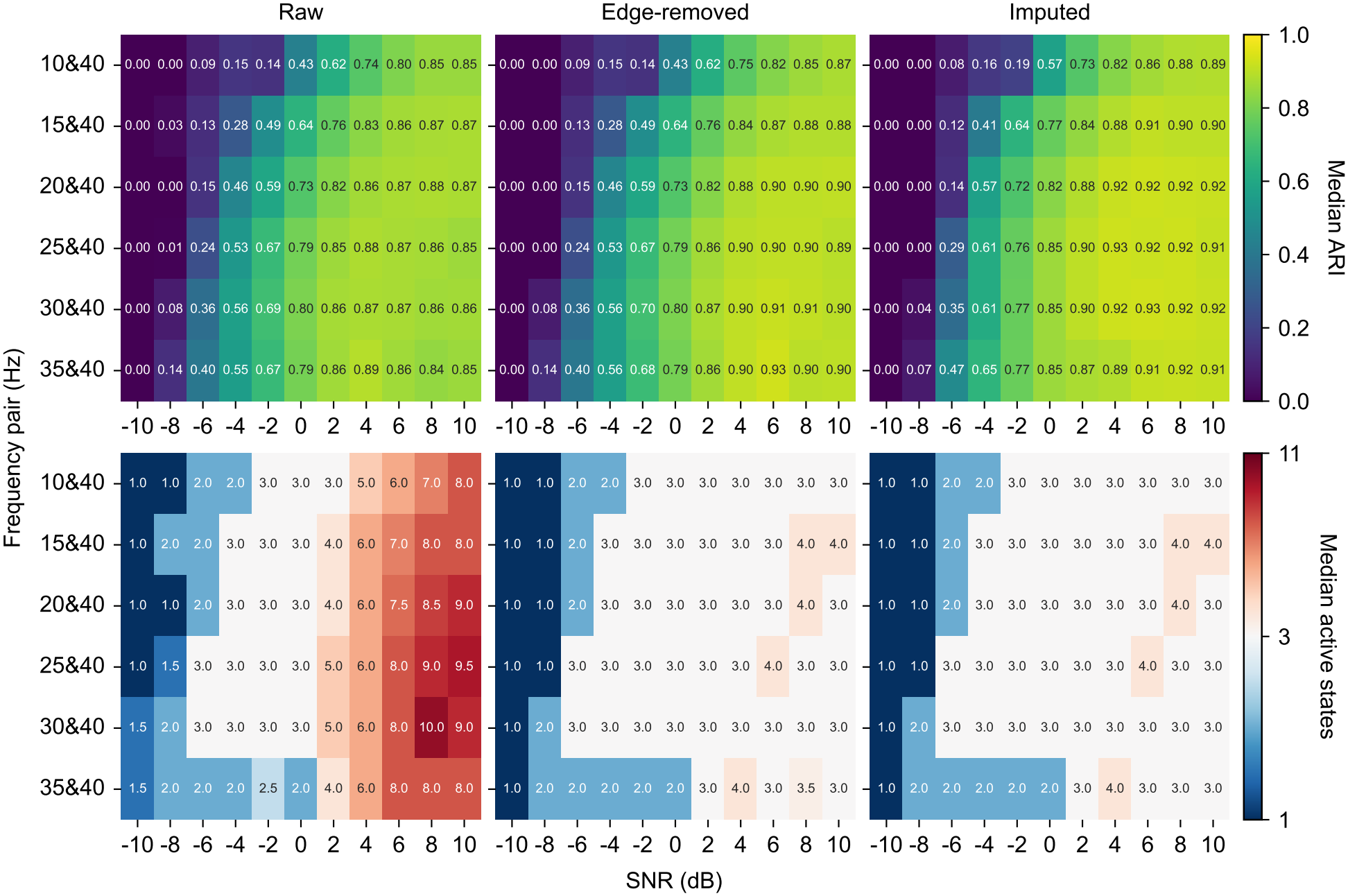
DP-GMM clustering accuracy and active-state counts before and after post-processing. Heatmaps show the median adjusted Rand index (ARI; top row) and median number of active states (bottom row) across 20 independent simulations containing two oscillatory states and one pink-noise state. Results are shown for six oscillatory-frequency pairs and SNRs from − 10 to 10 dB. Columns show the raw DP-GMM results after excluding components with negligible mixture weights, results after additional CV-based edge-state removal, and results after edge-state removal followed by *k*-NN imputation. Numbers within cells indicate the medians. The state-count color scale is centered on the true number of states, *K* = 3.

**Fig. S10.**
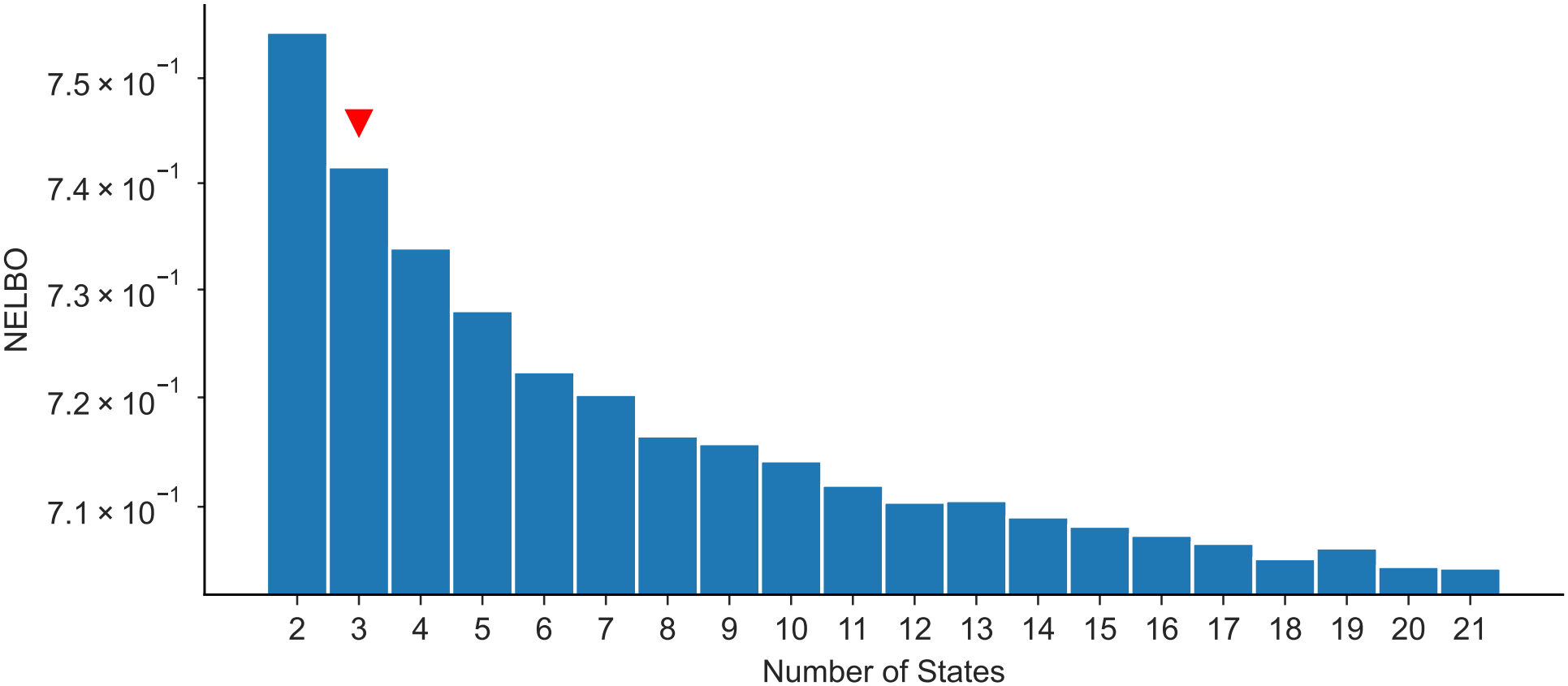
NELBO does not provide a clear criterion for selecting the true number of HMM states. HMM model selection by negative ELBO (NELBO) as a function of the number of states *K* ∈ {2, …, 21} on the three-state synthetic dataset. The red triangle marks the true state count, *K*_true_ = 3. The NELBO yielded no clear elbow or optimum at *K*_true_ = 3.

**Fig. S11.**
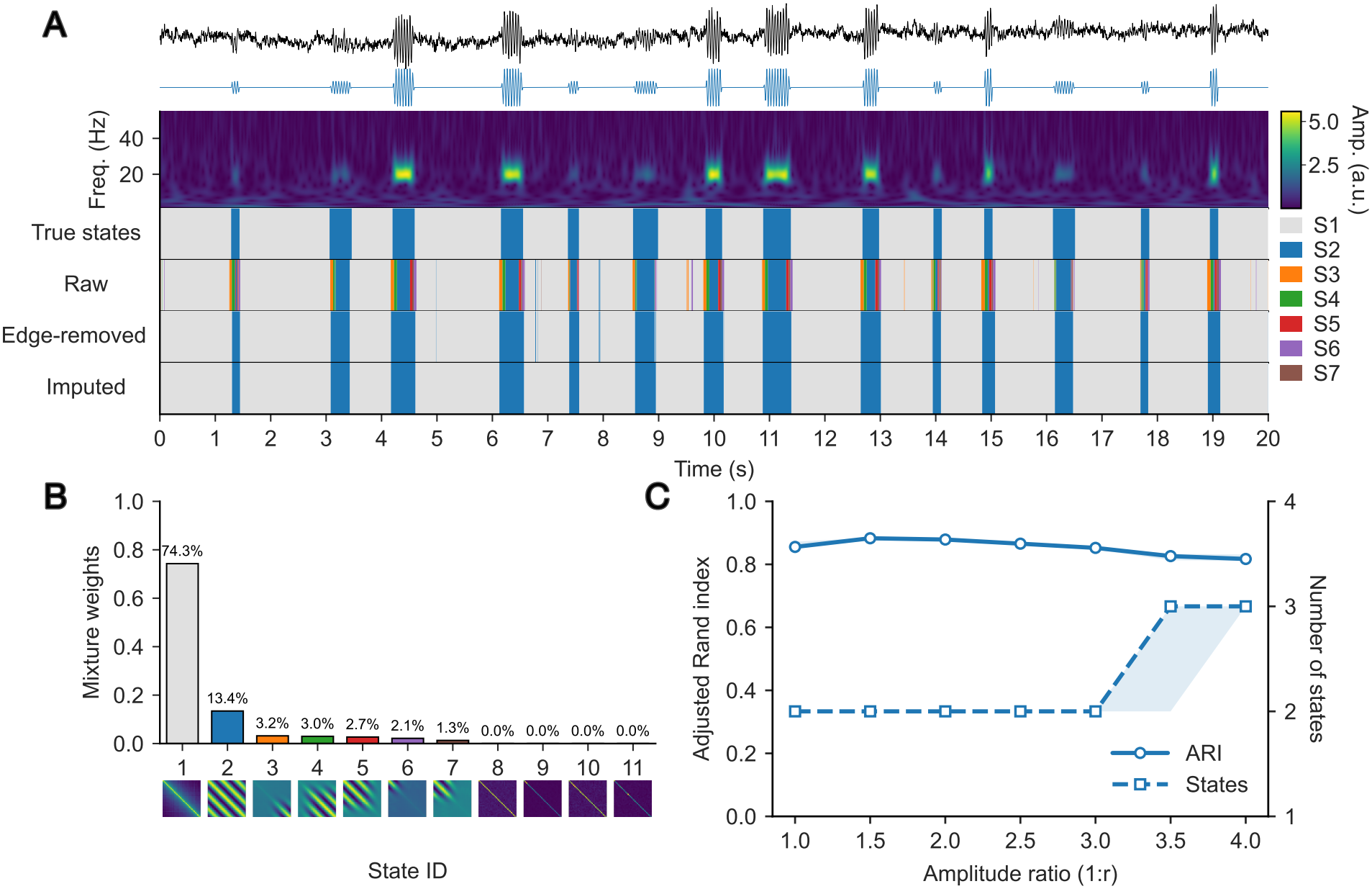
DP-GMM state identification under varying burst amplitudes. (A) Example signal containing 20 Hz oscillatory bursts with an amplitude ratio of 1:3. The lower-amplitude bursts were set to an SNR of 2 dB. From top to bottom: the simulated signal, oscillatory component, wavelet amplitude, ground-truth state sequence, and DP-GMM state sequences before edge-state removal, after edge-state removal, and after subsequent *k*-NN imputation. Both amplitude levels belong to the same ground-truth oscillatory state. (B) Posterior mixture weights and corresponding autocovariance matrices for all 11 candidate components before edge-state removal. (C) Median ARI (solid line with circles; left axis) and median number of active states (dashed line with squares; right axis) after edge-state removal and imputation, as a function of the amplitude ratio 1:*r*, with *r* ranging from 1 to 4 in increments of 0.5. Shading indicates the interquartile range across 20 independent 3-min simulations per condition, sampled at 250 Hz.

**Fig. S12.**
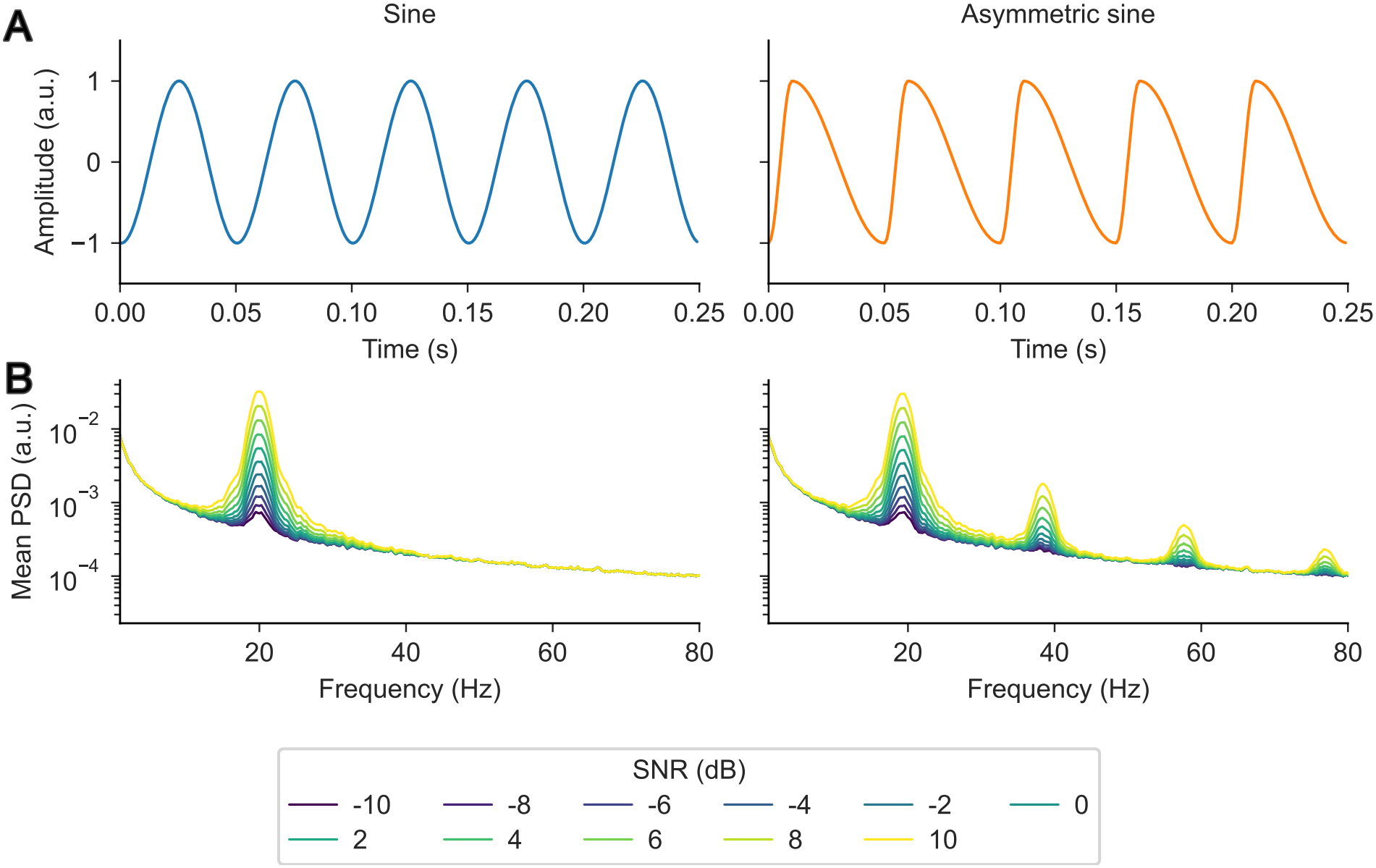
Simulated non-sinusoidal oscillations. (A) Example 20 Hz sine and asymmetric waves. The asymmetric wave has a rise–decay symmetry parameter of 0.2, meaning that the rising phase occupies only 20% of each cycle (i.e., a faster rise and slower decay). (B) Power spectral densities (PSDs) averaged across 20 simulations at each SNR. The PSDs of the asymmetric waves show harmonics near 40, 60, and 80 Hz.

**Fig. S13.**
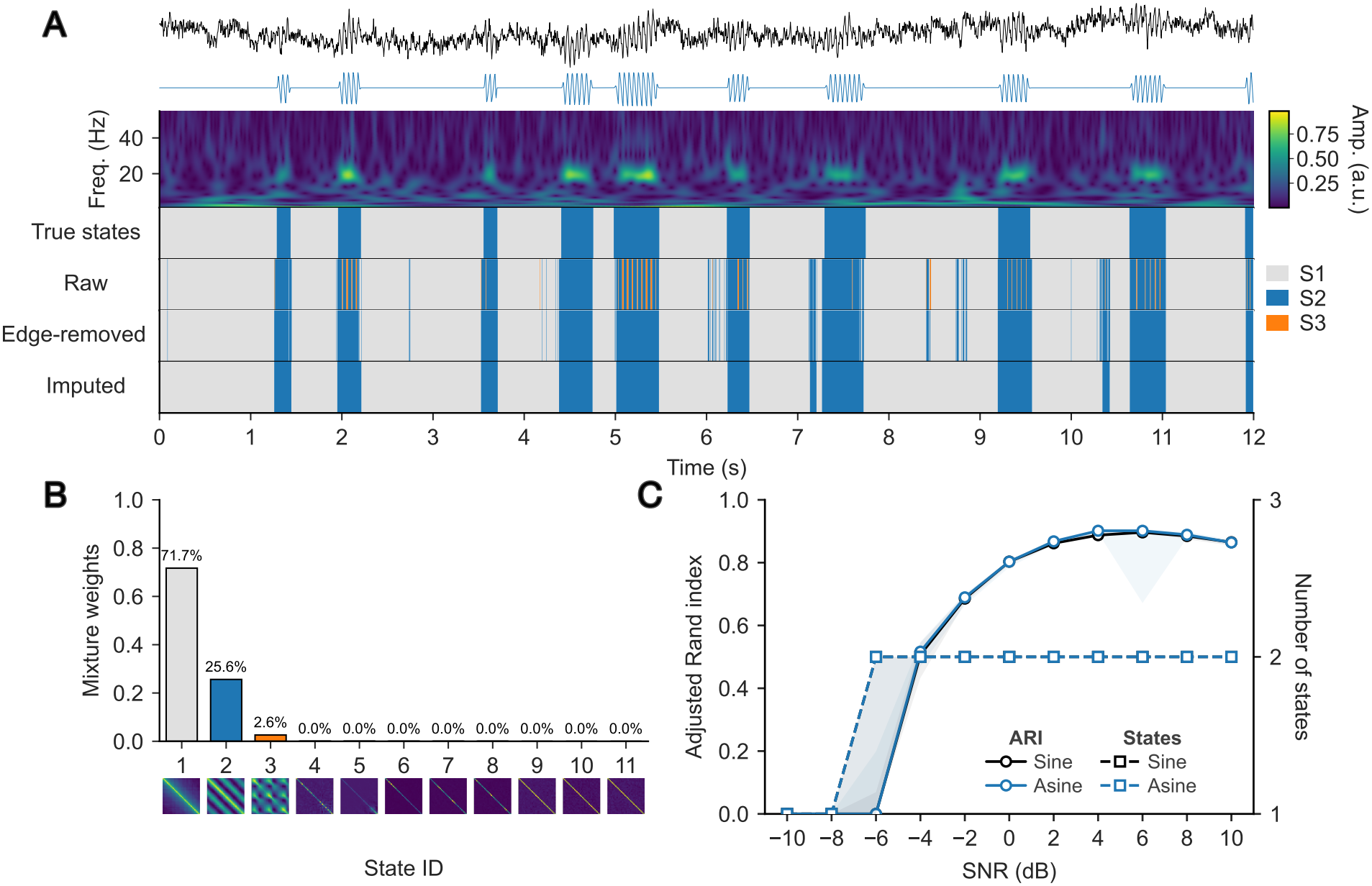
DP-GMM clustering of sinusoidal and asymmetric oscillatory bursts. (A) Example signal containing asymmetric 20 Hz bursts at an SNR of 2 dB. From top to bottom: the simulated signal, oscillatory component, wavelet amplitude, ground-truth state sequence, and DP-GMM state sequences before edge-state removal, after edge-state removal, and after subsequent *k*-NN imputation. (B) Posterior mixture weights and corresponding autocovariance matrices for all 11 candidate components before edge-state removal. (C) Median ARI (solid lines with circles; left axis) and median number of active states (dashed lines with squares; right axis) as a function of SNR, comparing sinusoidal bursts (black; Sine) and asymmetric bursts (blue; Asine) after edge-state removal and imputation. Shading indicates the interquartile range across 20 independent 3-min simulations per condition, sampled at 250 Hz.

**Fig. S14.**
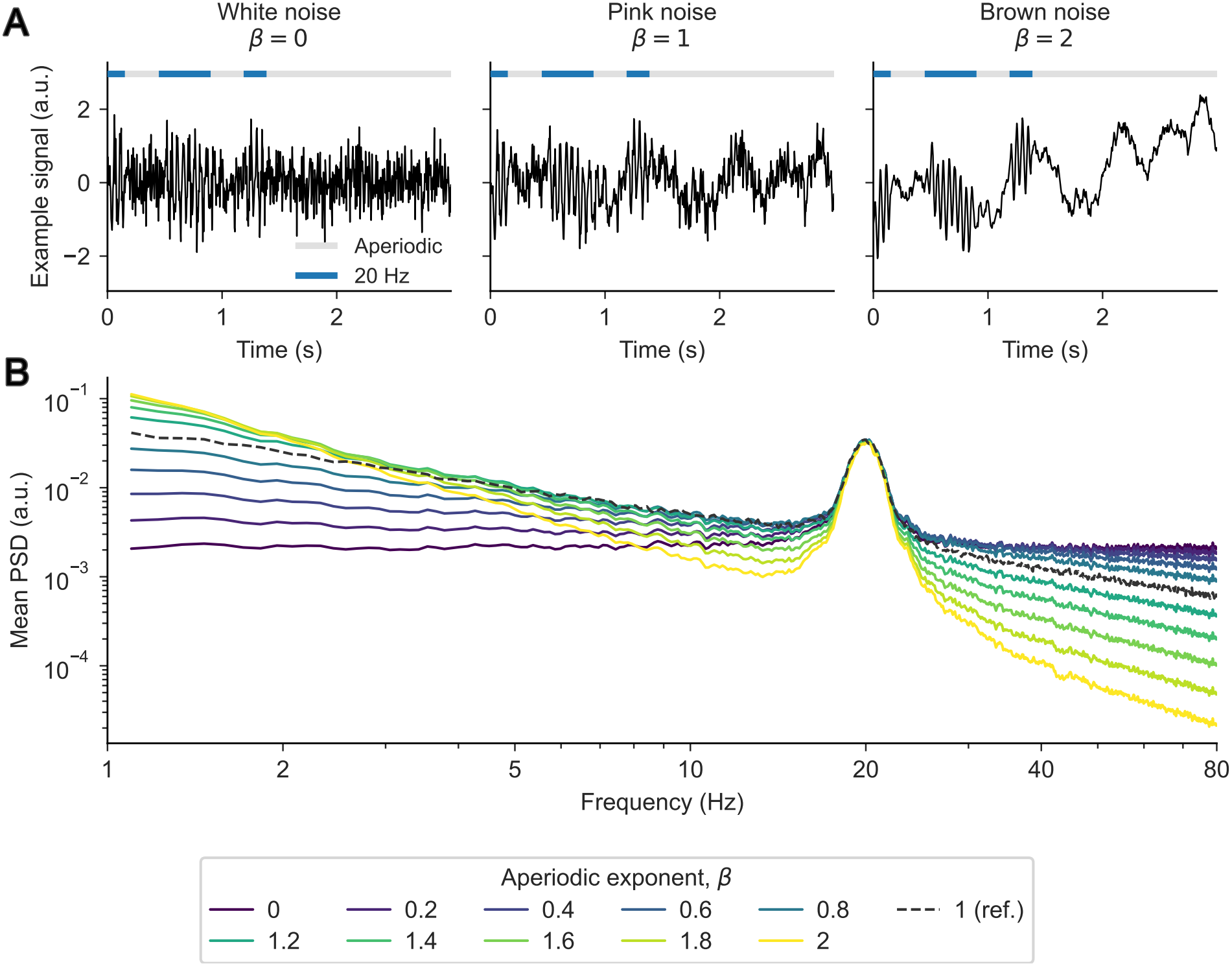
Synthetic signals and power spectra across aperiodic exponents. (A) Three-second examples of signals containing 20-Hz oscillatory bursts embedded in white (*β* = 0), pink (*β* = 1), or brown (*β* = 2) noise at a target SNR of 2 dB. Bars above each signal trace indicate periods containing only the aperiodic background (gray) and oscillatory periods (blue). (B) Mean PSDs for the varying aperiodic exponent *β*_2_, from 0 to 2 in increments of 0.2, excluding 1. The dashed gray curve shows the *β*_1_ = 1 reference. PSDs were estimated using only the blocks corresponding to *β*_2_ (or *β*_1_ = 1 for the baseline) and averaged across 20 simulations.

**Fig. S15.**
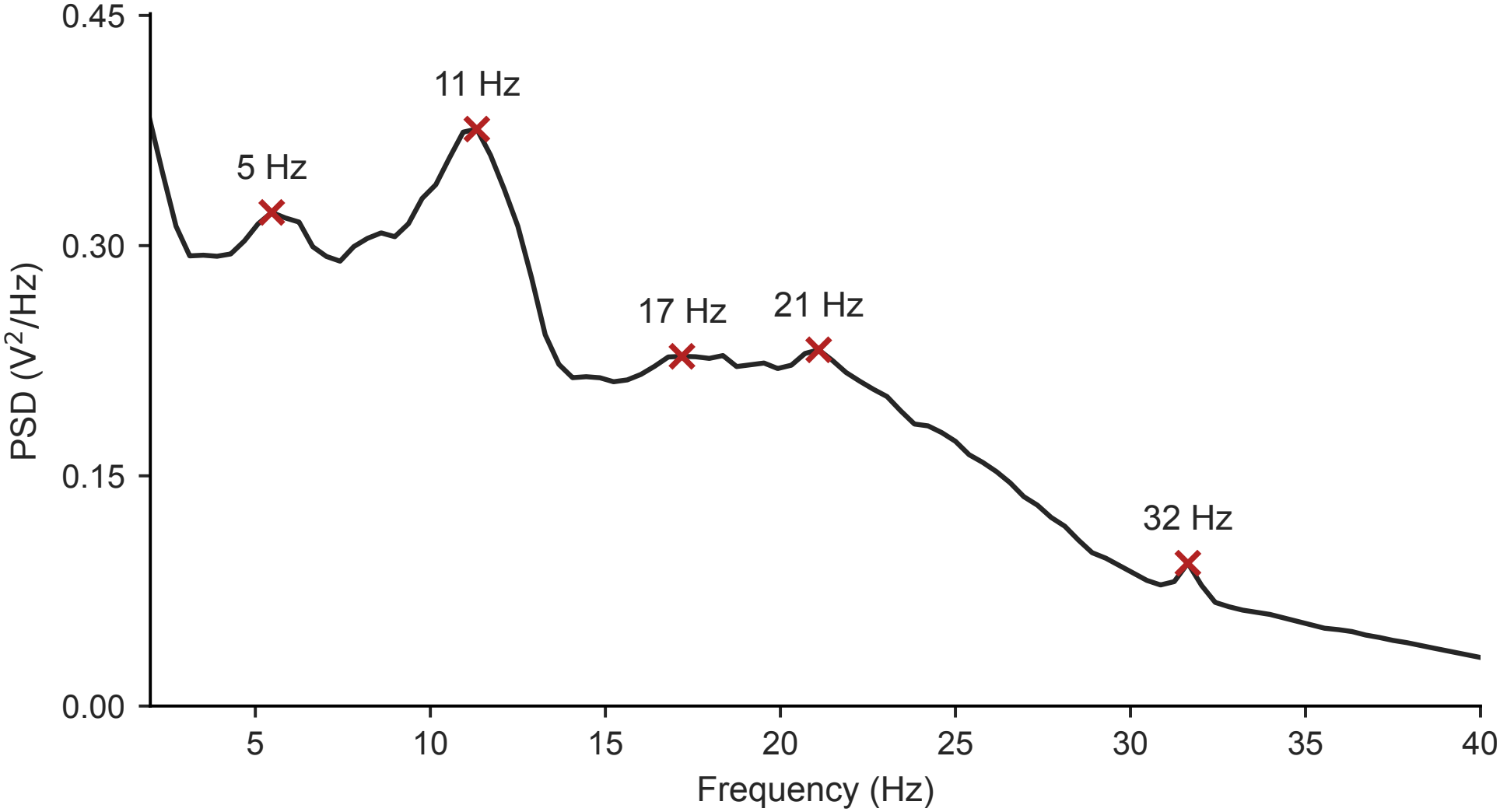
Candidate peak frequencies from the group-averaged resting-state PSD used to set the TDE and post-processing windows. Group-averaged power spectral density (PSD; V^2^/Hz) of CTF resting-state MEG, estimated with Welch’s method at *f*_*s*_ = 100 Hz. Local maxima (red crosses) were detected using a minimum peak separation of 4 Hz. The lowest-frequency peak at ∼ 2 Hz was excluded as likely reflecting the aperiodic 1*/f* background. The remaining candidate frequencies were 5, 11, 17, 21, and 32 Hz. Their mean (≈ 17 Hz) set the TDE window to 2.5 cycles (*E* = 15 lags; ≈ 0.145 s).

**Fig. S16.**
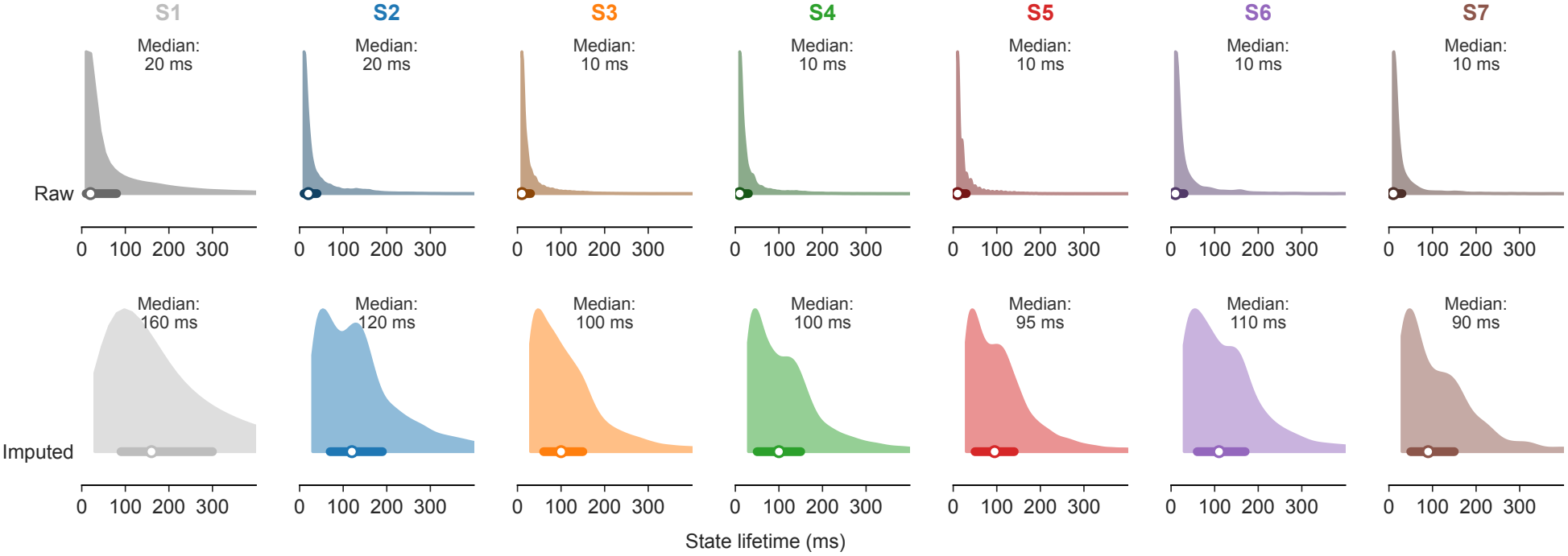
Brief state segments dominated lifetime distributions and were suppressed by *k*-NN imputation. State lifetimes were calculated from the decoded state sequences of all participants before *k*-NN imputation (top; raw) and after imputation (bottom; imputed). Bars indicate the median and interquartile range for each state. For visualization, only state lifetimes up to 400 ms are shown.

**Fig. S17.**
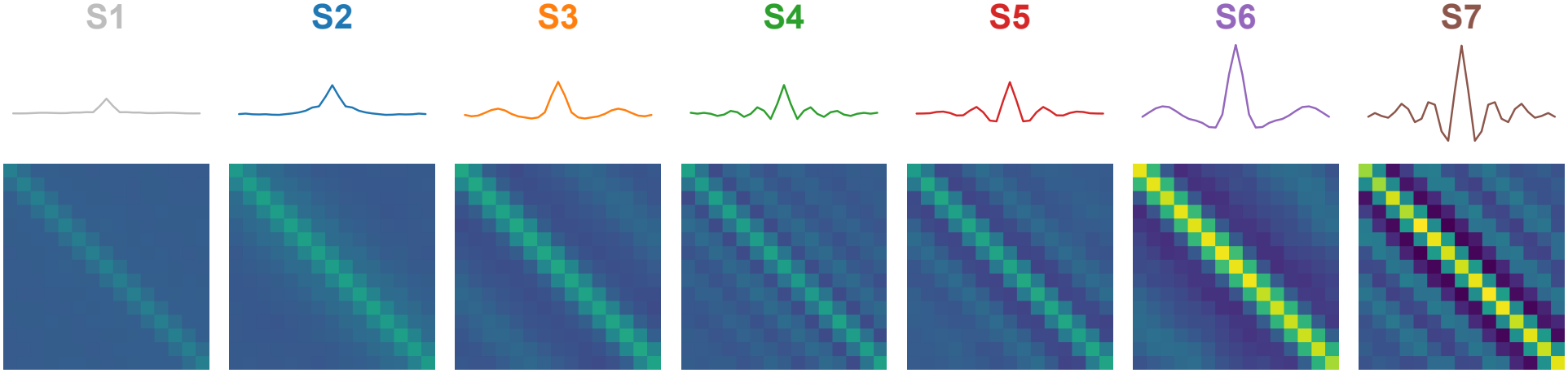
Estimated state-specific autocovariance matrices and corresponding autocovariance functions. Autocovariance matrices (bottom) and their corresponding autocovariance functions (top) are shown for the seven active states. Each autocovariance function was obtained by averaging matrix entries along diagonals corresponding to the same temporal lag. The matrices share a common color scale, and the autocovariance functions share a common vertical scale.

**Fig. S18.**
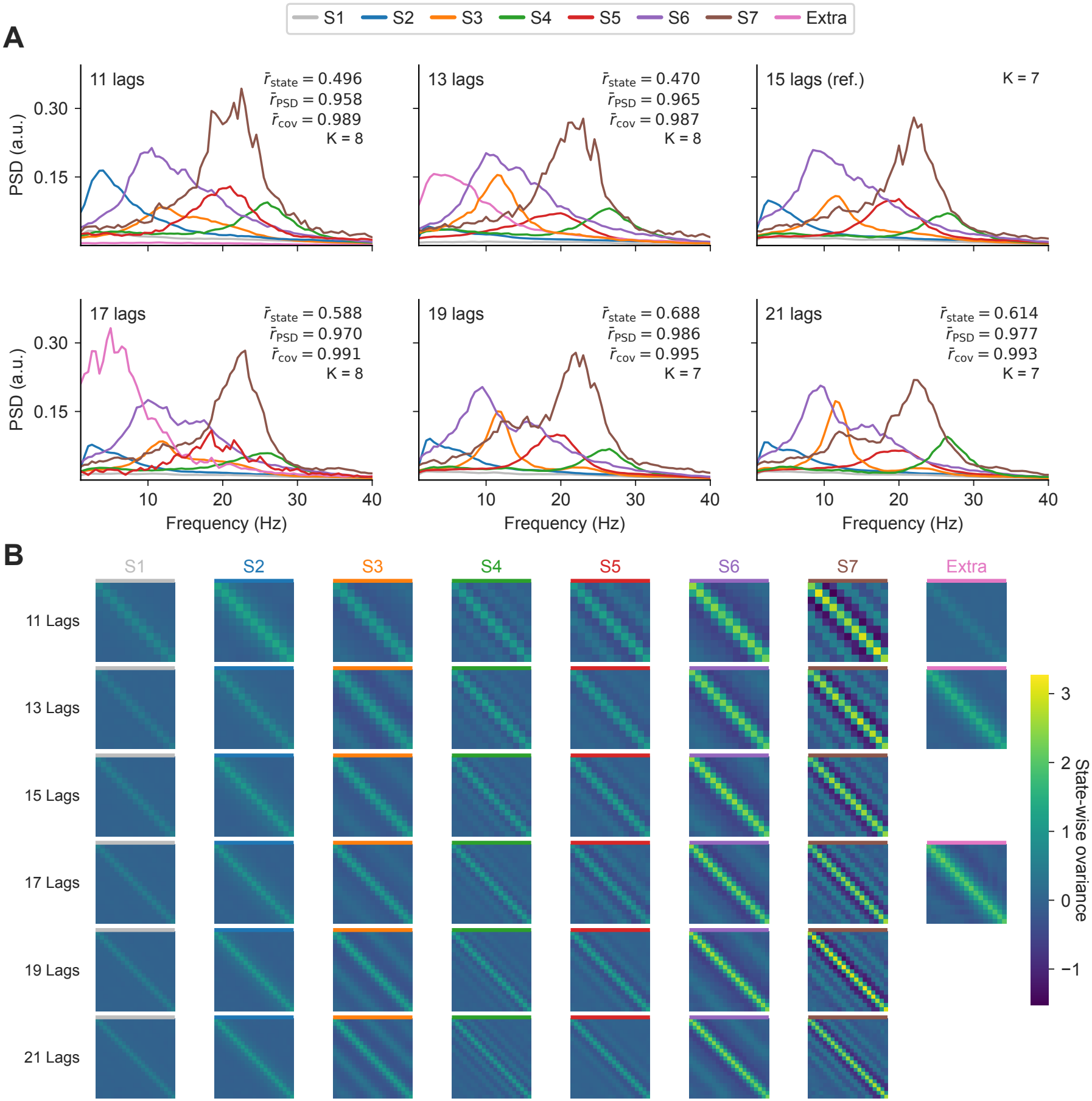
Sensitivity analysis of the DP-GMM to the number of embedding lags on the MEG dataset. Results are shown for 11, 13, 15, 17, 19, and 21 lags following edge-state removal and *k*-NN imputation with a 30 ms window. The 15-lag model was used as the reference. States were matched to the seven reference states through an optimal one-to-one assignment that maximized the sum of Pearson correlations between binary decoded-state time courses. “Extra” denotes an unmatched state. (A) State-specific PSDs. *K* denotes the number of active states. Annotations report mean Pearson correlations of matched decoded-state time courses 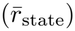, PSDs over 1–40 Hz 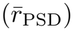, and covariance patterns 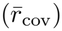. Before computing Pearson correlations, we converted the estimated categorical state sequence into a binary vector for each state. We averaged each of the three correlation measures across the same set of matched state pairs, excluding unmatched states. (B) Estimated state covariance matrices across embedding dimensions. Covariance matrices of different dimensions were displayed in equally sized square panels. To calculate 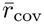, the centered 11 × 11 submatrix, corresponding to lag coordinates − 5 through +5, was extracted from each covariance matrix, including those of the reference model. These submatrices were flattened into 121-element vectors, and Pearson correlations were calculated for each matched state pair.

**Fig. S19.**
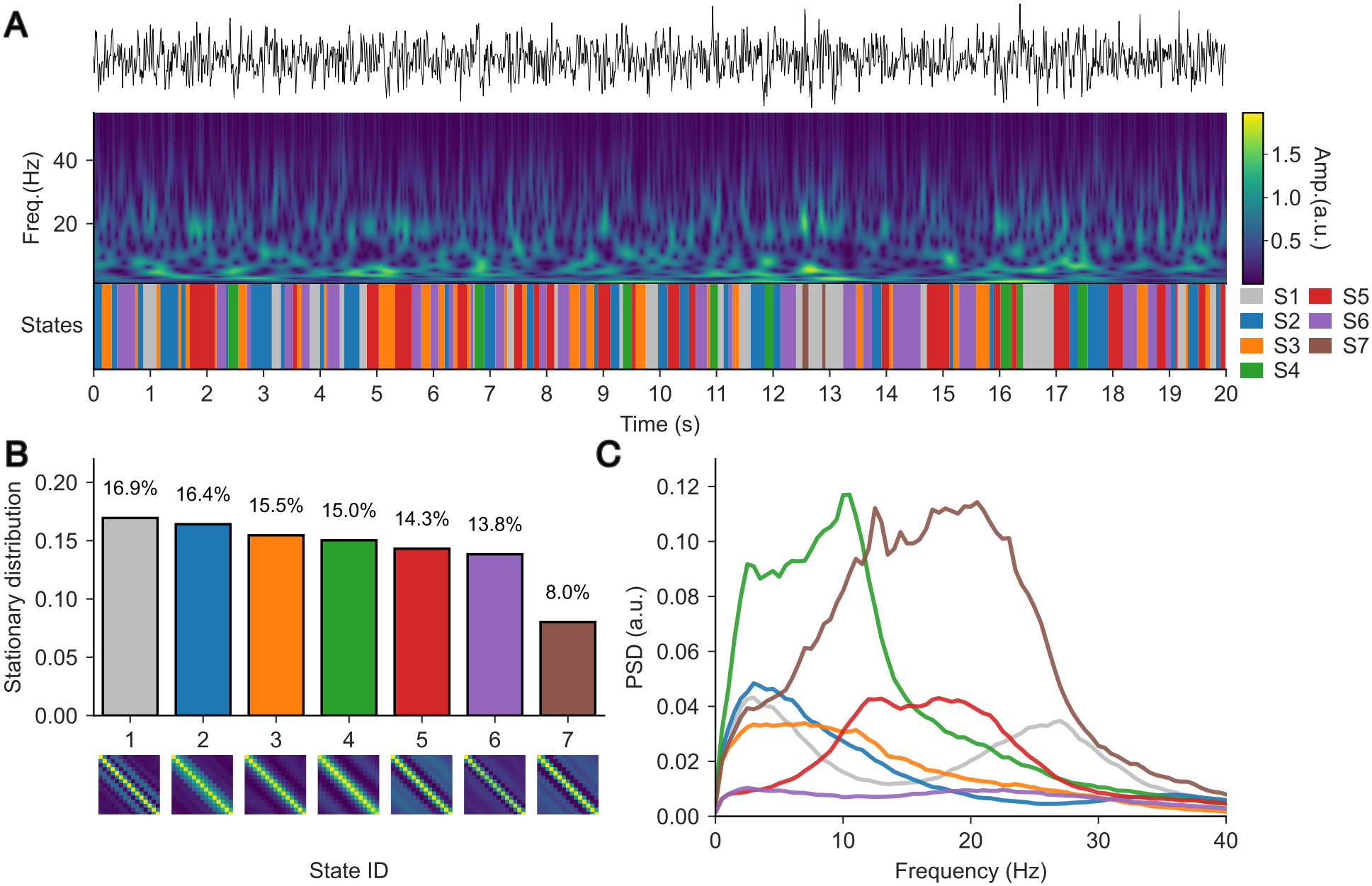
Oscillatory states were less clearly separated by the seven-state HMM. (A) Example CTF resting-state trace, wavelet representation, and decoded HMM state sequence (Viterbi path). (B) Estimated stationary state occupancies for *K* = 7, with state-specific autocovariance matrices. (C) State-wise power spectra. No post-hoc *k*-NN imputation was applied.

**Fig. S20.**
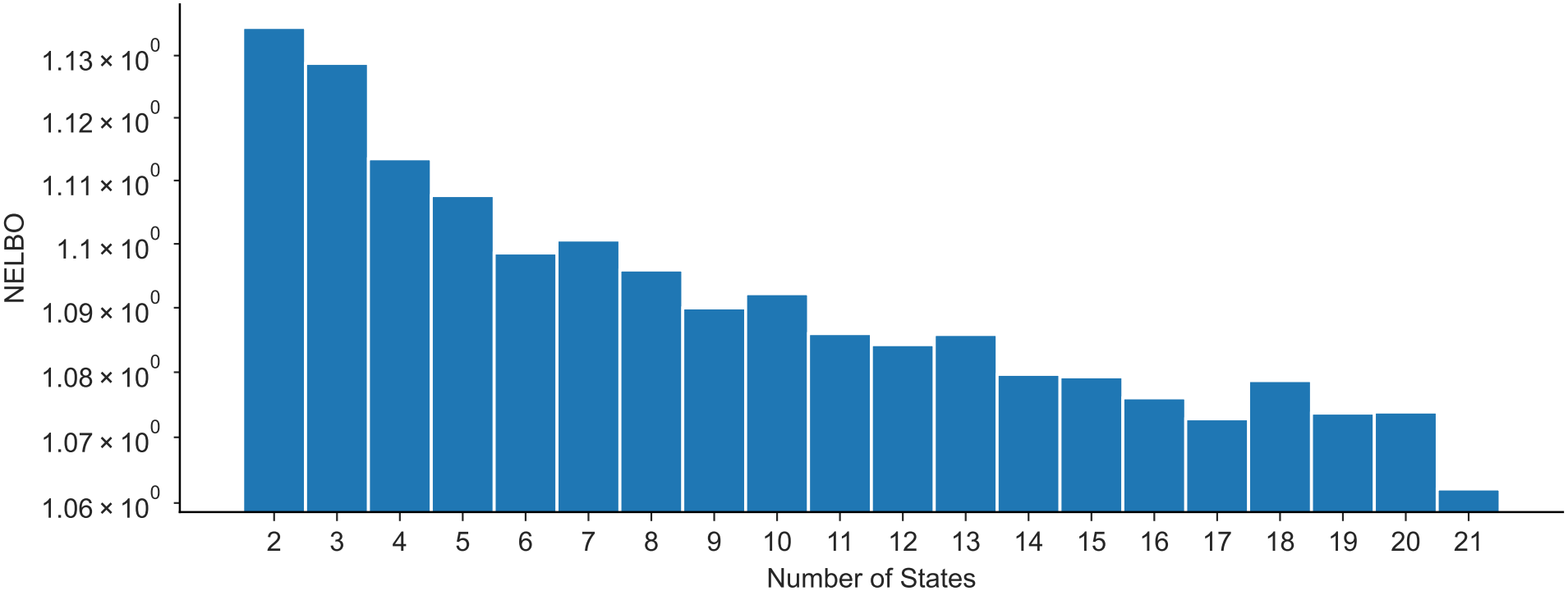
NELBO favors models with more states on real MEG data and does not yield a clear choice of *K*. Negative ELBO (NELBO) as a function of the number of states *K* ∈ { 2, …, 21} for HMMs fit to the CTF resting-state dataset. The NELBO decreases overall as *K* increases, providing no clear optimum at a small *K*.

**Fig. S21.**
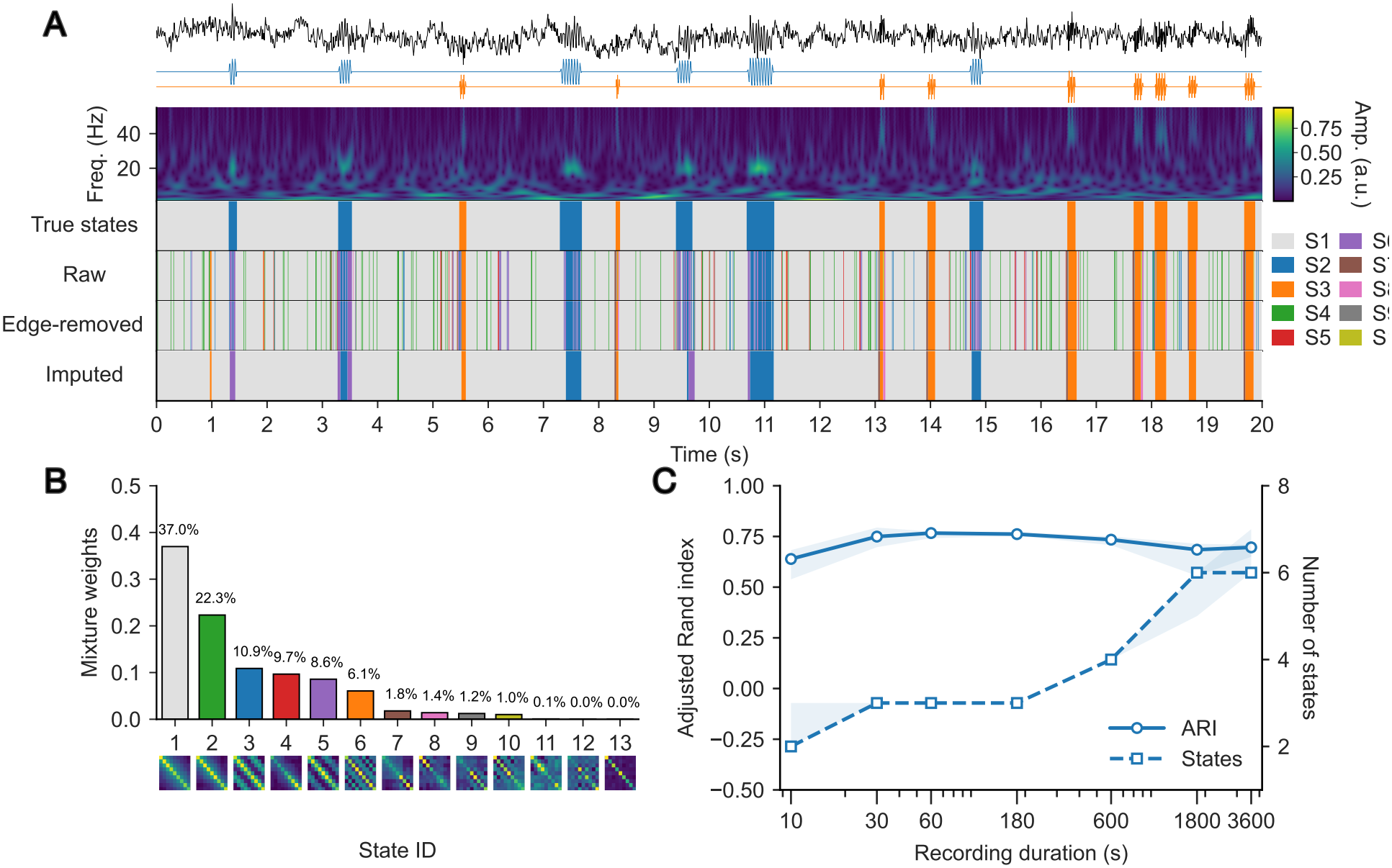
DP-GMM clustering performance and state-count estimates across recording durations. Twenty independent synthetic recordings contained 20 Hz and 40 Hz oscillatory bursts in pink noise at an SNR of 0 dB and a sampling rate of 100 Hz. Within each simulation, shorter recordings were taken from the beginning of the same 3600-s signal. The DP-GMM used nine embedding dimensions and an upper state bound of *K*_upper_ = ⌈ln *N* ⌉, where *N* is the number of samples in the recording. (A) A 20-s excerpt from a representative 3600-s recording, with the signal, oscillatory components, wavelet amplitude, ground-truth states, and decoded DP-GMM states before post-processing (raw), after edge-state removal, and after subsequent *k*-NN imputation. (B) Posterior mixture weights and corresponding autocovariance matrices for all 13 candidate components in the representative model. (C) Median adjusted Rand index (solid line, circles; left axis) and number of active states (dashed line, squares; right axis) after edge-state removal and *k*-NN imputation, across recording durations of 10, 30, 60, 180, 600, 1800, and 3600 s. Shaded bands indicate interquartile ranges across the 20 simulations. The recording-duration axis is logarithmic.

**Fig. S22.**
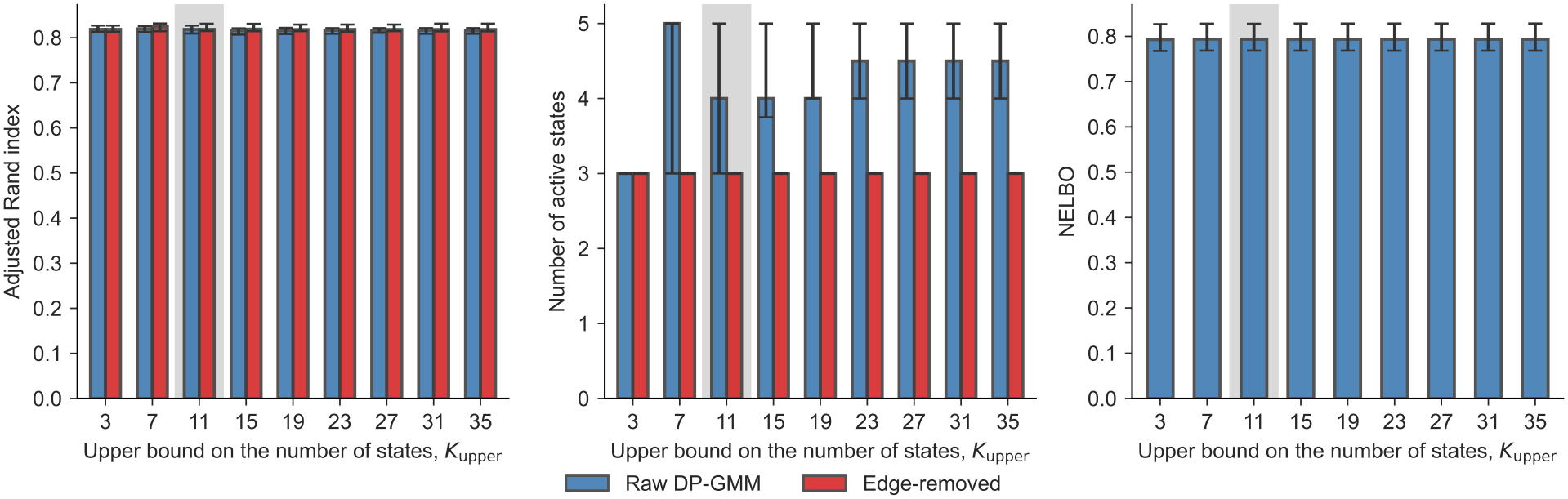
Sensitivity to the upper bound on the number of states. We varied *K*_upper_ from 3 to 35 across 20 three-minute simulated recordings sampled at 250 Hz. Each recording contained three ground-truth states corresponding to 20-Hz bursts, 40-Hz bursts, and pink noise alone. The bursts were embedded in pink noise at an SNR of 2 dB. The panels show the adjusted Rand index (left), the number of active states (middle), and the negative evidence lower bound (right). Clustering accuracy varied little across the tested values of *K*_upper_. The raw DP-GMM often identified more than three active states. Edge states accounted for this excess, rather than differences in *K*_upper_. After CV-based edge-state removal, the median number of active states was three at every tested upper bound. Blue and red bars show results before and after edge-state removal, respectively. Gray shading marks *K*_upper_ = 11.

## References

1. Varela F, Lachaux JP, Rodriguez E, Martinerie J. The brainweb: Phase synchronization and large-scale integration. Nature Reviews Neuroscience. 2001 Apr;2(4):229–39. Available from: https://www.nature.com/articles/35067550. doi:10.1038/35067550.

2. Buzsáki G, Draguhn A. Neuronal Oscillations in Cortical Networks. Science. 2004 Jun;304(5679):1926–9. Available from: https://www.science.org/doi/10.1126/science.1099745. doi:10.1126/science.1099745.

3. Siegel M, Donner TH, Engel AK. Spectral fingerprints of large-scale neuronal interactions. Nature Reviews Neuroscience. 2012 Feb;13(2):121–34. Available from: https://www.nature.com/articles/nrn3137. doi:10.1038/nrn3137.

4. Fries P. Rhythms for Cognition: Communication through Coherence. Neuron. 2015 Oct;88(1):220–35. Available from: https://linkinghub.elsevier.com/retrieve/pii/S0896627315008235. doi:10.1016/j.neuron.2015.09.034.

5. Cavanagh JF, Frank MJ. Frontal theta as a mechanism for cognitive control. Trends in cognitive sciences. 2014 Aug;18(8):414–21. Available from: https://pmc.ncbi.nlm.nih.gov/articles/PMC4112145/. doi:10.1016/j.tics.2014.04.012.

6. Schmidt R, Herrojo Ruiz M, Kilavik BE, Lundqvist M, Starr PA, Aron AR. Beta Oscillations in Working Memory, Executive Control of Movement and Thought, and Sensorimotor Function. J Neurosci. 2019 Oct;39(42):8231–8. doi:10.1523/JNEUROSCI.1163-19.2019.

7. Diekelmann S, Born J. The memory function of sleep. Nature Reviews Neuroscience. 2010 Feb;11(2):114–26. doi:10.1038/nrn2762.

8. Rasch B, Born J. About sleep’s role in memory. Physiological Reviews. 2013 Apr;93(2):681–766. Available from: https://www.physiology.org/doi/10.1152/physrev.00032.2012. doi:10.1152/physrev.00032.2012.

9. Zich C, Quinn AJ, Mardell LC, Ward NS, Bestmann S. Dissecting Transient Burst Events. Trends in Cognitive Sciences. 2020 Oct;24(10):784–8. Available from: https://www.ncbi.nlm.nih.gov/pmc/articles/PMC7653675/. doi:10.1016/j.tics.2020.07.004.

10. Muralidharan V, Aron AR, Schmidt R. Transient beta modulates decision thresholds during human action-stopping. NeuroImage. 2022 Jul;254:119145. Available from: https://www.sciencedirect.com/science/article/pii/S1053811922002725. doi:10.1016/j.neuroimage.2022.119145.

11. Muralidharan V, Aron AR, Cohen MX, Schmidt R. Two modes of midfrontal theta suggest a role in conflict and error processing. NeuroImage. 2023 Jun;273:120107. Available from: https://www.sciencedirect.com/science/article/pii/S1053811923002537. doi:10.1016/j.neuroimage.2023.120107.

12. Schmidt R, Rose J, Muralidharan V. Transient oscillations as computations for cognition: Analysis, modeling and function. Current Opinion in Neurobiology. 2023 Dec;83:102796. Available from: https://linkinghub.elsevier.com/retrieve/pii/S0959438823001216. doi:10.1016/j.conb.2023.102796.

13. Rayson H, Moreau Q, Gailhard S, Szul MJ, Bonaiuto JJ. Beta Burst Waveform Diversity: A Window onto Cortical Computation. The Neuroscientist. 2026 Feb;32(1):56–71. Available from: https://journals.sagepub.com/doi/10.1177/10738584251390779. doi:10.1177/10738584251390779.

14. Donoghue T, Schaworonkow N, Voytek B. Methodological considerations for studying neural oscillations. European Journal of Neuroscience. 2022;55(11-12):3502–27. Available from: https://onlinelibrary.wiley.com/doi/abs/10.1111/ejn.15361. doi:10.1111/ejn.15361.

15. Williams AH, Poole B, Maheswaranathan N, Dhawale AK, Fisher T, Wilson CD, et al. Discovering Precise Temporal Patterns in Large-Scale Neural Recordings through Robust and Interpretable Time Warping. Neuron. 2020 Jan;105(2):246–59.e8. Available from: https://www.cell.com/neuron/abstract/S0896-6273(19)30894-3. doi:10.1016/j.neuron.2019.10.020.

16. van Ede F, Quinn AJ, Woolrich MW, Nobre AC. Neural Oscillations: Sustained Rhythms or Transient Burst-Events? Trends in Neurosciences. 2018 Jul;41(7):415–7. Available from: https://www.ncbi.nlm.nih.gov/pmc/articles/PMC6024376/. doi:10.1016/j.tins.2018.04.004.

17. Shin H, Law R, Tsutsui S, Moore CI, Jones SR. The rate of transient beta frequency events predicts behavior across tasks and species. eLife. 2017 Nov;6:e29086. Available from: https://elifesciences.org/articles/29086. doi:10.7554/eLife.29086.

18. Little S, Bonaiuto J, Barnes G, Bestmann S. Human motor cortical beta bursts relate to movement planning and response errors. PLOS Biology. 2019 Oct;17(10):e3000479. Available from: https://dx.plos.org/10.1371/journal.pbio.3000479. doi:10.1371/journal.pbio.3000479.

19. Duchet B, Ghezzi F, Weerasinghe G, Tinkhauser G, Kühn AA, Brown P, et al. Average beta burst duration profiles provide a signature of dynamical changes between the ON and OFF medication states in Parkinson’s disease. PLOS Computational Biology. 2021 Jul;17(7):e1009116. Available from: https://journals.plos.org/ploscompbiol/article?id=10.1371/journal.pcbi.1009116. doi:10.1371/journal.pcbi.1009116.

20. Rayson H, Debnath R, Alavizadeh S, Fox N, Ferrari PF, Bonaiuto JJ. Detection and analysis of cortical beta bursts in developmental EEG Data. Developmental Cognitive Neuroscience. 2022 Apr;54:101069. Available from: https://pmc.ncbi.nlm.nih.gov/articles/PMC8816670/. doi:10.1016/j.dcn.2022.101069.

21. Widmann A, Schröger E, Maess B. Digital filter design for electrophysiological data – a practical approach. Journal of Neuroscience Methods. 2015 Jul;250:34–46. Available from: https://www.sciencedirect.com/science/article/pii/S0165027014002866. doi:10.1016/j.jneumeth.2014.08.002.

22. Langford ZD, Wilson CRE. Simulations reveal that beta burst detection may inappropriately characterize the beta band. Journal of Neurophysiology. 2025 Jul;134(1):10–9. Available from: https://journals.physiology.org/doi/full/10.1152/jn.00125.2024. doi:10.1152/jn.00125.2024.

23. Quinn AJ, van Ede F, Brookes MJ, Heideman SG, Nowak M, Seedat ZA, et al. Unpacking Transient Event Dynamics in Electrophysiological Power Spectra. Brain Topography. 2019 Nov;32(6):1020–34. doi:10.1007/s10548-019-00745-5.

24. Gohil C, Huang R, Roberts E, Van Es MW, Quinn AJ, Vidaurre D, et al. osl-dynamics, a toolbox for modeling fast dynamic brain activity. eLife. 2024 Jan;12:RP91949. Available from: https://elifesciences.org/articles/91949. doi:10.7554/eLife.91949.

25. Buckby J, Wang T, Fletcher D, Zhuang J, Takeo A, Obara K. Finding the number of latent states in hidden Markov models using information criteria. Environmental and Ecological Statistics. 2023 Dec;30(4):797–825. Available from: https://link.springer.com/10.1007/s10651-023-00584-5. doi:10.1007/s10651-023-00584-5.

26. Akaike H. Information Theory and an Extension of the Maximum Likelihood Principle. In: Parzen E, Tanabe K, Kitagawa G, editors. Selected Papers of Hirotugu Akaike. Springer Series in Statistics. New York, NY: Springer New York; 1998. p. 199–213. Available from: http://link.springer.com/10.1007/978-1-4612-1694-0_15. doi:10.1007/978-1-4612-1694-015.

27. Schwarz G. Estimating the Dimension of a Model. The Annals of Statistics. 1978 Mar;6(2):461–4. Available from: https://projecteuclid.org/journals/annals-of-statistics/volume-6/issue-2/Estimating-the-Dimension-of-a-Model/10.1214/aos/1176344136.full. doi:10.1214/aos/1176344136.

28. Bacci S, Pandolfi S, Pennoni F. A comparison of some criteria for states selection in the latent Markov model for longitudinal data. Advances in Data Analysis and Classification. 2014 Jun;8(2):125–45. Available from: https://doi.org/10.1007/s11634-013-0154-2. doi:10.1007/s11634-013-0154-2.

29. Pohle J, Langrock R, van Beest FM, Schmidt NM. Selecting the Number of States in Hidden Markov Models: Pragmatic Solutions Illustrated Using Animal Movement. Journal of Agricultural, Biological and Environmental Statistics. 2017 Sep;22(3):270–93. Available from: https://doi.org/10.1007/s13253-017-0283-8. doi:10.1007/s13253-017-0283-8.

30. Quinn AJ, Vidaurre D, Abeysuriya R, Becker R, Nobre AC, Woolrich MW. Task-Evoked Dynamic Network Analysis Through Hidden Markov Modeling. Frontiers in Neuroscience. 2018 Aug;12:603. Available from: https://www.frontiersin.org/article/10.3389/fnins.2018.00603/full. doi:10.3389/fnins.2018.00603.

31. Huang R, Gohil C, Woolrich M. Modelling variability in functional brain networks using embeddings. Imaging Neuroscience. 2026 Apr;4:IMAG.a.1188. Available from: https://doi.org/10.1162/IMAG.a.1188. doi:10.1162/IMAG.a.1188.

32. Gershman SJ, Blei DM. A tutorial on Bayesian nonparametric models. Journal of Mathematical Psychology. 2012 Feb;56(1):1–12. Available from: https://linkinghub.elsevier.com/retrieve/pii/S002224961100071X. doi:10.1016/j.jmp.2011.08.004.

33. Cho S, Choi JH. A guide towards optimal detection of transient oscillatory bursts with unknown parameters. Journal of Neural Engineering. 2023 Aug;20(4):046007. Available from: https://iopscience.iop.org/article/10.1088/1741-2552/acdffd. doi:10.1088/1741-2552/acdffd.

34. Hubert L, Arabie P. Comparing partitions. Journal of Classification. 1985 Dec;2(1):193–218. Available from: http://link.springer.com/10.1007/BF01908075. doi:10.1007/BF01908075.

35. Gates AJ, Ahn YY. The Impact of Random Models on Clustering Similarity. Journal of Machine Learning Research. 2017 Jul;18(87):1–28. Available from: https://www.jmlr.org/papers/v18/17-039.html.

36. Vinh NX, Epps J, Bailey J. Information theoretic measures for clusterings comparison: is a correction for chance necessary? In: Proceedings of the 26th Annual International Conference on Machine Learning. ICML ‘09. New York, NY, USA: Association for Computing Machinery; 2009. p. 1073–80. Available from: https://dl.acm.org/doi/10.1145/1553374.1553511. doi:10.1145/1553374.1553511.

37. Steinley D. Properties of the Hubert–Arabie Adjusted Rand Index. Psychological Methods. 2004;9(3):386–96. Available from: https://doi.apa.org/doi/10.1037/1082-989X.9.3.386. doi:10.1037/1082-989X.9.3.386.

38. Ferguson TS. A Bayesian Analysis of Some Nonparametric Problems. The Annals of Statistics. 1973 Mar;1(2):209–30. Available from: https://projecteuclid.org/journals/annals-of-statistics/volume-1/issue-2/A-Bayesian-Analysis-of-Some-Nonparametric-Problems/10.1214/aos/1176342360.full. doi:10.1214/aos/1176342360.

39. Ishwaran H, James LF. Gibbs Sampling Methods for Stick-Breaking Priors. Journal of the American Statistical Association. 2001 Mar;96(453):161–73. Available from: http://www.tandfonline.com/doi/abs/10.1198/016214501750332758. doi:10.1198/016214501750332758.

40. Barnard J, McCulloch R, Meng XL. Modeling covariance matrices in terms of standard deviations and correlations, with application to shrinkage. Statistica Sinica. 2000 Oct;10(4):1281–311. Available from: https://www3.stat.sinica.edu.tw/statistica/j10n4/j10n416/j10n416.htm.

41. Lewandowski D, Kurowicka D, Joe H. Generating random correlation matrices based on vines and extended onion method. J Multivar Anal. 2009 Oct;100(9):1989–2001. Available from: https://doi.org/10.1016/j.jmva.2009.04.008. doi:10.1016/j.jmva.2009.04.008.

42. Korwar RM, Hollander M. Contributions to the Theory of Dirichlet Processes. The Annals of Probability. 1973 Aug;1(4):705–11. Available from: https://projecteuclid.org/journals/annals-of-probability/volume-1/issue-4/Contributions-to-the-Theory-of-Dirichlet-Processes/10.1214/aop/1176996898.full. doi:10.1214/aop/1176996898.

43. Rabiner LR. A tutorial on hidden Markov models and selected applications in speech recognition. Proceedings of the IEEE. 1989 Feb;77(2):257–86. Available from: http://ieeexplore.ieee.org/document/18626/. doi:10.1109/5.18626.

44. Fraser AM. Hidden Markov models and dynamical systems. Philadelphia, Pa: Society for Industrial and Applied Mathematics; 2008. Available from: https://doi.org/10.1137/1.9780898717747. doi:10.1137/1.9780898717747.

45. Viterbi A. Error bounds for convolutional codes and an asymptotically optimum decoding algorithm. IEEE Transactions on Information Theory. 1967 Apr;13(2):260–9. Available from: https://ieeexplore.ieee.org/document/1054010. doi:10.1109/TIT.1967.1054010.

46. Linderman SW, Chang P, Harper-Donnelly G, Kara A, Li X, Duran-Martin G, et al. Dynamax: A Python package for probabilistic state space modeling with JAX. Journal of Open Source Software. 2025 Apr;10(108):7069. Available from: https://joss.theoj.org/papers/10.21105/joss.07069. doi:10.21105/joss.07069.

47. Wingate D, Weber T. Automated Variational Inference in Probabilistic Programming; 2013. ArXiv:1301.1299 [stat]. Available from: http://arxiv.org/abs/1301.1299. doi:10.48550/arXiv.1301.1299.

48. Ranganath R, Gerrish S, Blei D. Black Box Variational Inference. In: Kaski S, Corander J, editors. Proceedings of the Seventeenth International Conference on Artificial Intelligence and Statistics. vol. 33 of Proceedings of Machine Learning Research. PMLR; 2014. p. 814–22. Available from: https://proceedings.mlr.press/v33/ranganath14.html.

49. Hoffman MD, Blei DM, Wang C, Paisley J. Stochastic Variational Inference. Journal of Machine Learning Research. 2013 May;14(40):1303–47. Available from: https://www.jmlr.org/papers/v14/hoffman13a.html.

50. Blei DM, Kucukelbir A, McAuliffe JD. Variational Inference: A Review for Statisticians. Journal of the American Statistical Association. 2017 Apr;112(518):859–77. Available from: http://arxiv.org/abs/1601.00670. doi:10.1080/01621459.2017.1285773.

51. Phan D, Pradhan N, Jankowiak M. Composable Effects for Flexible and Accelerated Probabilistic Programming in NumPyro; 2019. ArXiv:1912.11554 [stat]. Available from: http://arxiv.org/abs/1912.11554. doi:10.48550/arXiv.1912.11554.

52. Bingham E, Chen JP, Jankowiak M, Obermeyer F, Pradhan N, Karaletsos T, et al. Pyro: Deep Universal Probabilistic Programming. Journal of Machine Learning Research. 2019 Feb;20(28):1–6. Available from: https://www.jmlr.org/papers/v20/18-403.html.

53. Obermeyer F, Bingham E, Jankowiak M, Chiu J, Pradhan N, Rush AM, et al. Tensor Variable Elimination for Plated Factor Graphs. In: Chaudhuri K, Salakhutdinov R, editors. Proceedings of the 36th International Conference on Machine Learning. vol. 97 of Proceedings of Machine Learning Research. PMLR; 2019. p. 4871–80. Available from: https://proceedings.mlr.press/v97/obermeyer19a.html.

54. Schulman J, Heess N, Weber T, Abbeel P. Gradient Estimation Using Stochastic Computation Graphs. In: Advances in Neural Information Processing Systems. vol. 28. Curran Associates, Inc.; 2015. p. 3528–36. Available from: https://proceedings.neurips.cc/paper_files/paper/2015/hash/de03beffeed9da5f3639a621bcab5dd4-Abstract.html.

55. Cole S, Donoghue T, Gao R, Voytek B. NeuroDSP: A package for neural digital signal processing. Journal of Open Source Software. 2019 Apr;4(36):1272. Available from: https://joss.theoj.org/papers/10.21105/joss.01272. doi:10.21105/joss.01272.

56. Feingold J, Gibson DJ, DePasquale B, Graybiel AM. Bursts of beta oscillation differentiate postperformance activity in the striatum and motor cortex of monkeys performing movement tasks. Proceedings of the National Academy of Sciences. 2015 Nov;112(44):13687–92. Available from: https://www.pnas.org/doi/full/10.1073/pnas.1517629112. doi:10.1073/pnas.1517629112.

57. MEG UK Scientific Research Community. The scientific community for MEG research in the United Kingdom and Ireland [Internet]. MEG UKI; n.d. [cited 2025 Mar 15]. Available from: https://meguk.ac.uk/.

58. Gao R, Peterson EJ, Voytek B. Inferring synaptic excitation/inhibition balance from field potentials. NeuroImage. 2017 Sep;158:70–8. Available from: https://www.sciencedirect.com/science/article/pii/S1053811917305621. doi:10.1016/j.neuroimage.2017.06.078.

59. Ostlund B, Donoghue T, Anaya B, Gunther KE, Karalunas SL, Voytek B, et al. Spectral parameterization for studying neurodevelopment: How and why. Developmental Cognitive Neuroscience. 2022 Apr;54:101073. Available from: https://linkinghub.elsevier.com/retrieve/pii/S1878929322000172. doi:10.1016/j.dcn.2022.101073.

60. Donoghue T. A Systematic Review of Aperiodic Neural Activity in Clinical Investigations. European Journal of Neuroscience. 2025 Oct;62(7):e70255. Available from: https://doi.org/10.1111/ejn.70255. doi:10.1111/ejn.70255.

61. Wilson LE, Da Silva Castanheira J, Baillet S. Time-resolved parameterization of aperiodic and periodic brain activity. eLife. 2022 Sep;11:e77348. Available from: https://elifesciences.org/articles/77348. doi:10.7554/eLife.77348.

62. Cho H, Adamek M, Willie JT, Brunner P. Novel cyclic homogeneous oscillation detection method for high accuracy and specific characterization of neural dynamics. eLife. 2024 Sep;12:RP91605. Available from: https://doi.org/10.7554/eLife.91605.3. doi:10.7554/eLife.91605.3.

63. Cole S, Voytek B. Cycle-by-cycle analysis of neural oscillations. Journal of Neurophysiology. 2019 Aug;122(2):849–61. Available from: https://journals.physiology.org/doi/full/10.1152/jn.00273.2019. doi:10.1152/jn.00273.2019.

64. Cole SR, Voytek B. Brain Oscillations and the Importance of Waveform Shape. Trends in Cognitive Sciences. 2017 Feb;21(2):137–49. Available from: https://linkinghub.elsevier.com/retrieve/pii/S1364661316302182. doi:10.1016/j.tics.2016.12.008.

65. Belluscio MA, Mizuseki K, Schmidt R, Kempter R, Buzsáki G. Cross-Frequency Phase–Phase Coupling between Theta and Gamma Oscillations in the Hippocampus. Journal of Neuroscience. 2012 Jan;32(2):423–35. Available from: https://www.jneurosci.org/content/32/2/423. doi:10.1523/JNEUROSCI.4122-11.2012.

66. de Cheveigné A, Nelken I. Filters: When, Why, and How (Not) to Use Them. Neuron. 2019 Apr;102(2):280–93. Available from: https://www.sciencedirect.com/science/article/pii/S0896627319301746. doi:10.1016/j.neuron.2019.02.039.

67. Haegens S, Cousijn H, Wallis G, Harrison PJ, Nobre AC. Inter- and intra-individual variability in alpha peak frequency. NeuroImage. 2014 May;92:46–55. doi:10.1016/j.neuroimage.2014.01.049.

68. Mahjoory K, Schoffelen JM, Keitel A, Gross J. The frequency gradient of human resting-state brain oscillations follows cortical hierarchies. eLife. 2020 Aug;9:e53715. Available from: https://doi.org/10.7554/eLife.53715. doi:10.7554/eLife.53715.

69. Yu SZ. Hidden semi-Markov models. Artificial Intelligence. 2010 Feb;174(2):215–43. Available from: https://linkinghub.elsevier.com/retrieve/pii/S0004370209001416. doi:10.1016/j.artint.2009.11.011.

70. Beal MJ, Ghahramani Z, Rasmussen CE. The Infinite Hidden Markov Model. In: Advances in Neural Information Processing Systems. vol. 14. MIT Press; 2001. p. 577–84. Available from: https://papers.nips.cc/paper_files/paper/2001/hash/e3408432c1a48a52fb6c74d926b38886-Abstract.html.

71. Johnson MJ, Willsky AS. Bayesian nonparametric hidden semi-Markov models. Journal of Machine Learning Research. 2013 Feb;14(20):673–701. Available from: https://www.jmlr.org/papers/v14/johnson13a.html.

72. Tan E, Algar S, Corrêa D, Small M, Stemler T, Walker D. Selecting embedding delays: An overview of embedding techniques and a new method using persistent homology. Chaos: An Interdisciplinary Journal of Nonlinear Science. 2023 Mar;33(3):032101. Available from: https://pubs.aip.org/cha/article/33/3/032101/2881154/Selecting-embedding-delays-An-overview-of. doi:10.1063/5.0137223.

73. Liu AA, Henin S, Abbaspoor S, Bragin A, Buffalo EA, Farrell JS, et al. A consensus statement on detection of hippocampal sharp wave ripples and differentiation from other fast oscillations. Nature Communications. 2022 Oct;13(1):6000. Available from: https://www.nature.com/articles/s41467-022-33536-x. doi:10.1038/s41467-022-33536-x.

74. Synigal SR, Teoh ES, Lalor EC. Including Measures of High Gamma Power Can Improve the Decoding of Natural Speech From EEG. Frontiers in Human Neuroscience. 2020 Apr;14:130. Available from: https://pmc.ncbi.nlm.nih.gov/articles/PMC7200998/. doi:10.3389/fnhum.2020.00130.

75. Attar ET. Review of electroencephalography signals approaches for mental stress assessment. Neurosciences. 2022 Oct;27(4):209–15. Available from: https://pmc.ncbi.nlm.nih.gov/articles/PMC9749579/. doi:10.17712/nsj.2022.4.20220025.

76. Staresina BP, Bergmann TO, Bonnefond M, van der Meij R, Jensen O, Deuker L, et al. Hierarchical nesting of slow oscillations, spindles and ripples in the human hippocampus during sleep. Nature Neuroscience. 2015 Nov;18(11):1679–86. Available from: https://www.nature.com/articles/nn.4119. doi:10.1038/nn.4119.

77. Ouedraogo DW, Lenck-Santini PP, Marti G, Robbe D, Crépel V, Epsztein J. Abnormal UP/DOWN Membrane Potential Dynamics Coupled with the Neocortical Slow Oscillation in Dentate Granule Cells during the Latent Phase of Temporal Lobe Epilepsy. eNeuro. 2016 May;3(3):ENEURO.0017-16.2016. Available from: https://www.eneuro.org/lookup/doi/10.1523/ENEURO.0017-16.2016. doi:10.1523/ENEURO.0017-16.2016.

